# Food web structure alters ecological communities top-heaviness with little effect on the biodiversity-functioning relationship

**DOI:** 10.1101/2020.06.10.144949

**Authors:** Eva Delmas, Daniel B. Stouffer, Timothée Poisot

## Abstract

In a rapidly changing world, the composition, diversity and structure of ecological communities face many threats. Biodiversity-Ecosystem Functioning (BEF) and community food-chain analyses have focused on investigating the consequences of these changes on ecosystem processes and the resulting functions. These different and diverging conceptual frameworks have each produced important results and identified a set of important mechanisms, that shape ecosystem functions. But the disconnection between these frameworks, and the various simplifications of the study systems are not representative of the complexity of real-world communities. Here we use food webs as a more realistic depiction of communities, and use a bioenergetic model to simulate their biomass dynamics and quantify the resulting flows and stocks of biomass. We use tools from food web analysis to investigate how the predictions from BEF and food-chain analyses fit together, how they correlate to food-web structure and how it might help us understand the interplay between various drivers of ecosystem functioning. We show that food web structure is correlated to the community’s efficiency in storing the captured biomass, which may explain the distribution of biomass (top heaviness) across the different trophic compartments (producers, primary and secondary consumers). While we know that ecological network structure is important in shaping ecosystem dynamics, identifying structural attributes important in shaping ecosystem processes and synthesizing how it affects various underpinning mechanisms may help prioritize key conservation targets to protect not only biodiversity but also its structure and the resulting services.

## 1 Introduction

Understanding the consequence of diversity on ecosystems process rates has become one of the ecologists priorities since we realized that human activities threaten both species existence (MEA 2005) and the structure of the interactions between them (Poisot et al. 2015) at a global scale. A variety of analyses have been conducted to investigate this particular problematic, and have produced important results (Tilman et al. 2014; Plas 2019) but the divide between them in concepts and approaches has hindered our ability to synthesize their results to build a general theoretical framework (Raffaelli 2006; Duffy et al. 2007; Hines et al. 2019).

In particular, while some have focused on understanding the effect of “horizontal diversity” (the diversity of species, usually within trophic levels) on ecosystems process rates (Biodiversity-Ecosystem functioning analyses or BEF) through mechanisms derived from competition (Loreau & Hector 2001; Tilman et al. 2014), conversely others have focused on the effect of “vertical diversity” (the number of trophic level, or food chain length) by aggregating species into trophic compartments along the food chain and using concepts from food-chain theory (Fretwell 1987; Duffy et al. 2007; Loreau 2010). Both approaches have produced important results, such as the paradigmatic BEF relationship (Tilman et al. 1996), or the consequences of trophic cascades on community structure (Paine 1980; Polis & Strong 1996; Borer et al. 2006; Gruner et al. 2008); but to build a mechanistic understanding of diversity effects on ecosystem processes rates in natural communities, comprising many species along the food chain, we need to be able to reconcile both dimensions of diversity, as well as the different approaches used to study their effect in a single framework (Ives et al. 2005; Duffy et al. 2007; Thompson et al. 2012; Barbier & Loreau 2019).

Recent efforts towards this goal have generally framed their analysis in food webs (Poisot et al. 2013; Schneider et al. 2016; Wang & Brose 2017; Wang et al. 2019; Buzhdygan et al. 2020). Studying the flux of biomass in ecological communities represented as their underlying network of trophic interaction – a map of whom eats whom, or food web – provide a powerful framework to investigate ecosystem functioning (Pascual & Dunne 2006; Thompson et al. 2012). Fluxes of nutrients and energy (commonly represented as biomass) in ecosystems represent the fundamental ecosystem process sustaining organisms, allowing their growth, underlying interaction between them and ultimately constraining the persistence and structure of the community (Barnes et al. 2018). As such, biomass fluxes represent a useful common currency to analyze, model and ultimately gain understanding of the aggregated ecosystem processes they underlay, from primary production to pressure of top carnivores (DeAngelis 1992), providing a way to estimate ecosystem multi-functionality (defined as the provisioning of multiple ecosystem functions; Barnes et al. 2018) and the potential to sustain various ecosystem services (Soliveres et al. 2016). By mapping trophic interactions, food webs also map biomass routes through the communities, the dynamics of the transfer of biomass through these routes can be modelled using adapted consumer-resource models such as Yodzis & Innes (1992). In parallel, the structure of these routes can be analyzed through a set of measures, adapted from graph theory, that carry a diversity of ecological information such as species degree of specialization, distance to producers, etc. (Dunne 2006; Delmas et al. 2019). This results in an ideal framework for modelling communities functioning, and identifying its drivers.

First results from theoretical analyses of the BEF relationship in food webs show that the positive - hitherto paradigmatic - relationship between the diversity of species in a community and functioning rates appears to be dependent on the structure of the network (Thébault & Loreau 2006). For example, networks containing few VS. many generalist species would have qualitatively different BEF relationships. More recent in-silico experiments that followed have usually framed their analyses in more realistic communities, i.e., containing more species and using models that generate food webs with realistic structures of interactions (such as the niche model; Williams & Martinez 2000). The results of these studies show that the dependence on structure of the BEF relationship may not be as strong as previously thought, there still seems to be a positive relationship between diversity and functioning rates (Schneider et al. 2016; Wang & Brose 2017; Wang et al. 2019), but this relationship is relatively noisy. These studies, along with other more theoretical work (Poisot et al. 2013), have offered mechanisms that could extend the action of selection and complementarity to a framework involving competition and consumption, but no consensus has yet been reached.

Most of these recent analyses, using food webs to investigate diversity effects, although they integrate both dimensions of diversity (vertical and horizontal), do not necessarily attempt to bridge the gap between the conceptual frameworks that have emerged from the analysis of their effects in isolation (namely BEF and trophic chain theories). The analysis of trophic chains dynamics in particular has produced important results for understanding the functioning of ecosystems (Ives et al. 2005). Among these results is the fact that changes in richness (and thus the functioning of a trophic level since the BEF relationship is valid within the different compartments; Cardinale et al. 2006a) are reflected, via the trophic cascade, on all the other trophic levels (Duffy et al. 2007). In parallel the same kind of approach shows that the type of trophic interaction can also vary the strength of the trophic cascade. For example, depending on the proportion of omnivores (or intra-guild predators) in a community, a variation in consumer richness will not have the same impact on plant biomass (losing a predator leads to a reduction in plant richness, whereas a decrease in the richness of omnivores leads to an increase in their biomass; Polis & Strong (1996)).

One important results from food chain theory, and more specifically from analyzing the effect of community-level trophic cascades (as opposed as species-level; Polis 1999), is the influence on community shapes. By shape we refer to what is generally called trophic structure, i.e., a qualification of the distribution of biomass or abundance along the different trophic levels of the community (cascade or pyramid, top or bottom heavy). We use the term shape here to avoid confusion with the food-web network structure. Recent studies by Barbier & Loreau (2019) and Galiana et al. (2020) present a framework to reconcile predictions resulting from the use of different approaches in the study of food chains (energetic vs. dynamic). Their results show that community shape can theoretically inform us on the strength and direction of trophic control (what they call donor control vs. antagonistic feedback). While these results are still theoretical, we know that top-heavy (TH) and bottom-heavy (BH) food webs for example do emerge under different energetic constrains reflecting different internal dynamic processes (Leroux & Loreau 2008; McCauley et al. 2018; Woodson et al. 2018). Similarly pyramidal or cascade shapes could also be a reflection of other energetic or dynamic constraints (Barbier & Loreau 2019; Galiana et al. 2020).

We believe that understanding the reciprocal feedback between biomass fluxes – underpinning ecosystem processes – food-web structure and community shape is an important key in the search for the mechanisms driving diversity effects on ecosystem functioning in complex communities. Investigating these relationship may help us understand the interdependence between the different drivers of ecosystem functioning, and in particular the way diversity and structure interact and drive ecosystem processes. We expect for example that the BEF relationship will be different depending on community shape, reflecting different energetic constraints on the dynamic transfers of biomass through trophic interactions. To test our hypothesis, we used food webs to represent communities, and simulated their biomass dynamics using a consumer-resource bioenergetic model adapted to food webs (Yodzis & Innes 1992; Williams et al. 2007). We investigated the potential link between the emerging shape of the communities, their food-web structure and BEF relationship. If our hypothesis proves to be valid, this would offer a possible explanation for the apparent idiosyncrasy of the BEF relationship in food webs. It would also provide ideas about the possible mechanisms responsible for the diversity-functioning relationship in complex ecological communities.

## 2 Methods

### 2.1 Biomass dynamics

To estimate biomass dynamics in food webs, we used a well-established bioenergetic food-web model (BEFWm). This model is an extension of Yodzis & Innes (1992) consumer-resource model to multiple resources and consumers (Williams et al. 2007). Biomass dynamics in this model hinge on three main dynamic processes described in the driving eq. 1 below: autotrophic production (first term of eq. 1), transfer of biomass through consumption (second and third terms of eq. 1), and loss of biomass because of metabolism (fourth term of eq. 1) and imperfect assimilation (e_ij_). The details of the model for plant growth (G_i_(N)), functional response (F_ji_) and all parameter values are described in supplementary material (appendix 1, section S1).

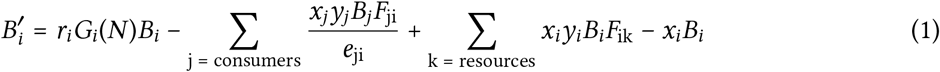

Plant growth through autotrophic production is the basic process that sustains communities. Competition between plants for abiotic resources is a fundamental process of ecosystem functioning (Loreau et al. 2001; Cardinale et al. 2006b). We made this competition explicit by combining a well established (Tilman 1982; Huisman & Weissing 1999) and empirically tested (Passarge et al. 2006) model of nutrient intake for producers with a food-web consumer-resource model (following Brose 2008). The resulting bioenergetic model integrates the basic mechanism from which BEF relationships emerge – competition – with mechanisms that result from consumption (e.g., transfer efficiency). This approach yields more realistic results with fewer extinctions than classical consumer-resource models (Brose 2008), as it integrates feedback between bottom-up transfer of biomass from plants to consumers and top-down control of plant biomass by consumers.

In the nutrient intake model (Brose et al. 2005b, 2005a; Brose 2008), all producers share two nutrients *N*_1_ and *N*_2_. Nutrients availability is determined by their respective rates of supply, turnover and consumption by plants. Plants all have different half-saturation density for both nutrients, which results in a hierarchy of competition between them, as a lower half-saturation means a higher intake efficiency. As plants do not need all nutrients in the same quantities, we set nutrient content in plants (c_1_ and c_2_ for respectively N_1_ and N_2_) to reflect this. We set c_1_ = 1 and c_2_ = 0.5, meaning that plants need a higher quantity of N_1_ than N_2_. The half-saturation for *N*_1_ is thus the primary driver of plants competitive hierarchy. The model equations are described in supplementary material (appendix 1, section S1.1.1).

Biomass is transferred through trophic interactions according to a multi-species functional response (Williams et al. 2007). This extension of the classical functional response accounts for consumers (resources) having multiple resources (consumers) and therefore accounts for both apparent and exploitative competition. We chose to implement a Holling type III functional response (Holling 1959; Real 1977) with homogeneous consumption among a consumer’s resources. Finally, biomass is lost through metabolism and imperfect assimilation. The functional response and its parameters are described in supplementary material (appendix 1, section S1.1.2).

The biological rates controlling these processes – namely the growth, maximum consumption and metabolic rates – are all dependant of two things: species metabolic class (vertebrate or invertebrate) and typical adult body size. In other words, we have an allometric scaling of biological rates with body size, with different allometric coefficient depending on the metabolic class.

### 2.2 Generating realistic food webs

We generated food webs with realistic structural properties using the Allometric Diet Breadth Model (ADBM, Petchey et al. 2008). This model is based on optimal foraging theory and the allometry of foraging variables with body size. As such, it needs to be initialized with a vector containing species typical adult body sizes. It predicts realistically interactions from empirical food webs (Petchey et al. 2008; but see Allesina 2011), and important food webs structural properties such as diet breadth and food web connectance, two potentially strong drivers of biomass dynamics.

To predict interactions between species, the ADBM model works in two steps. First it calculates the profitability (P_ij_, or rate of energy intake) for each pair of species in the community. Then, it selects the links that maximize it. Profitability is expressed as a function of the net energy gained through trophic interactions (which scales linearly with the consumer body size), the encounter rate (which depends on the density of the resource species here expressed as biomass and the allometrically scaled attack rate) and the handling time. We chose to implement the “ratio” method for estimating handling time as it is supposed to yield more accurate results (Petchey et al. 2008). In this formulation, handling time is estimated differently depending on the body-size ratio between a consumer and its potential resource. If the size different is too big (bigger than a chosen threshold; see section S2) then we assume that the focus consumer is not able to consume the focus resource. See section S2 and Petchey et al. (2008) for more detail on the model’s equations and parameter values.

### 2.3 Numerical experimental design

In order to generate realistic food webs that still express a generate wide range of richness and structure, we initialized the ADBM model with body-mass data from an empirical community: the Benguela pelagic community [Yodzis (1998); Brose et al. (2016); tab. S3 in appendix 1, supplementary material]. We chose this particular community because of the ability of the ADBM model to reproduce it quite well (Petchey et al. 2008). We sampled species from those present in this community and recorded their body mass and metabolic type (vertebrate or invertebrate). The body masses were used to generate food-web interaction matrices (see fig. 1, top left panel), and metabolic types were also used to calculate biological rates later used for the biomass dynamics simulations.

**Figure 1.**
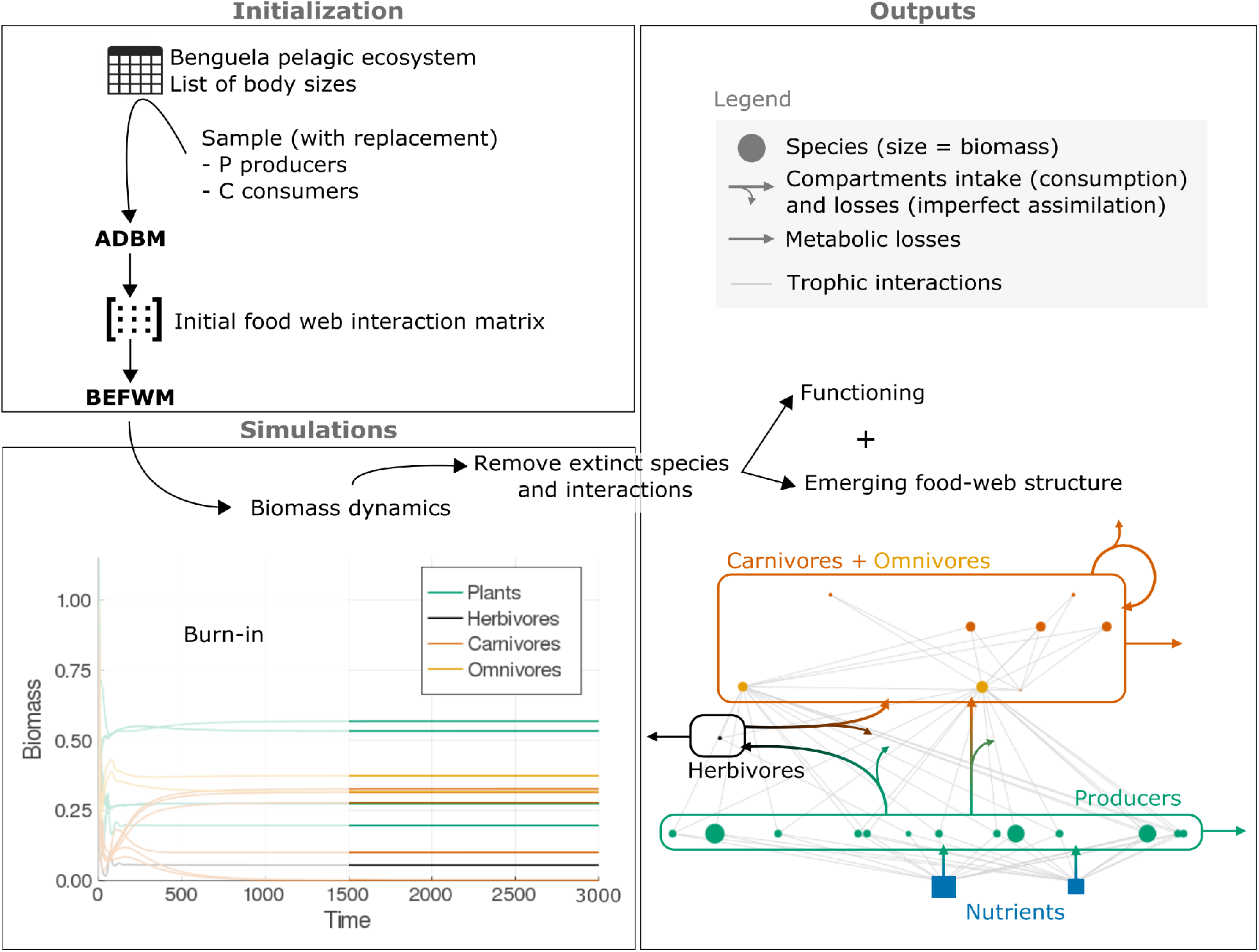
Conceptual representation of our experiments. Initialization (top left). We start by generating food webs with different levels of producer and consumer richness by sampling P producers and C consumers in a list of species (Benguela ecosystem as described in Yodzis, 1998). We retrieve the species body masses and metabolic classes. We pass the body masses to the Allometric Diet Breadth Model (ADBM, Petchey et al., 2008) to generate realistic food web interaction matrices. Simulation (bottom left). These matrices are passed to the Bioenergetic below 10 during the second half of the simulation are considered as extinct, all interactions from and to them are Food-Web model (Yodzis & Inès, 1992) to simulate their biomass dynamics. All species that have an average biomass below 10^−6^ during the second half of the simulation are considered as extinct, all interactions from and to them are removed. Outputs (right side). Emerging food-web structures, total biomasses and fluxes are calculated at the species level over the second half of the simulations and aggregated by trophic compartments (carnivores and omnivores in respectively dark and light orange, herbivores in black, producers in green). The whole system depends on the two basal nutrients (blue).

We generated food webs with 10 levels of producers richness (from 2 to 20), and 7 levels of consumer richness (from 5 to 35). For each of these 70 combinations of total species richness (7 to 55), we randomly generated up to 100 different food webs with the ADBM model. For each food web, we randomly drew the producers half-saturation density for the two shared nutrients, to generate a random hierarchy of competition among the different food webs. This design produces food webs with different richness, structure and ability to extract nutrients.

Because we wanted to set different typical consumer-resource body-mass ratio to explore the effect of allometric scaling, we did not use the original sampled body mass (used to generate food webs) in the simulations to calculate biological rates but reassigned body masses based on a sampled consumer-resource body-mass ratio (*Z*). A species body mass (M_*i*_) is calculated from *Z* and its trophic level (*t*_*i*_) in the following way: *M*_*i*_ = *Z*_*ti*_. Using this method provides us with a wider range of size structures. As allometric scaling is an important determinant of biomass dynamics and food web stability (Brose et al. 2006; Brose 2008), we believe that it is important to investigate its effect on functioning.

For each of these webs, we simulated biomass dynamics using the BioEnergeticFoodWeb package (v. 1.1.2; Delmas et al. 2020), a Julia (v. 1.3.1; Bezanson et al. 2017) implementation of the bioenergetic model as described above in the section Biomass dynamics. Biomass dynamics for each food web were simulated for each food web were simulated for 3000 time steps (see fig. 1, bottom left panel), as systems usually reach the dynamic equilibrium before 1000 time steps. Species extinctions were triggered when species biomass reach 10^−6^. These extinctions caused a new structure to emerge as all links from and to these species disappear. This emerging structure (see fig. 1 right panel) is the one that is later used for the analyses. Codes and data used for these analyses are available at https://osf.io/yuezq/?view_only=2378095a26d9414489fbcd4a72753a27.

For every food web that reached equilibrium with three compartments – food webs without secondary consumers were discarded – we estimated its functioning, structure and shape. Functioning here refers to the quantification of the fluxes and stocks within the food webs. Biomass fluxes (represented by arrows in fig. 1, right panel) in food webs is the basic process underpinning ecosystem functioning, and reflects many of the functioning typically studied in BEF analyses (Barnes et al. 2018). We thus extracted the biomass values at each time step for all species and calculated fluxes (which correspond here to species intakes, that is ∑_k = resources_ *x*_*i*_*y*_*i*_*B*_*i*_*F*_*ik*_ for consumers and *r*_*i*_*G*_*i*_(*N*)*B*_*i*_ for producers) and stocks (total biomass) at the species level using the equations of the model for corresponding process (see the corresponding sections in the sup. mat.). Both the fluxes and stocks were averaged over the second half of the iteration to compensate for potential oscillations. These quantities were later aggregated at the compartment level (plants, primary and secondary consumers) to generate results at the food chain scale. As a precision we define omnivores as species that can feed on both animals and producers.

To compare the biomass to intake relationships of communities with and without consumption, we also simulated the biomass dynamics of communities with producers only. As we needed to see a variation in intake, we simulated communities with varying richness (1 to 20) and supply rate (1 to 10). As intake is always close to its maximum value in the absence of consumption, we used a linear regression to extrapolate for a wider range of intake. Without consumption, the relationship between intake and stored biomass should indeed be linear (metabolic losses scale linearly with species biomass).

Structure refers to the organization of trophic interactions between species within the food webs. Once extinct species and the interactions to and from them were discarded, we measured the food web connectance, height (Dunne 2006) and motif profile (Milo 2002). Motifs are the different N-species sub-webs that can exist within a network (here *N* = 3). They are the simplest building blocks of networks. We focused here on 4 motifs that represent fundamental and widely studied trophic modules (omnivory or intraguild predation A → B → C ← A, food chains A → B → C, apparent A ← B → C and exploitative A → B ← C competition, note that in this paper arrows go from consumer to resource) and have been linked to food web dynamics (Stouffer et al. 2007; Borrelli 2015). We used the Julia package EcologicalNetworks.jl (v0.3.0; Poisot et al. 2020) and the python3 package pymfinder (Mora et al. 2018) to analyze food web structure.

Food web shape is defined here as what we usually call *trophic structure* in the literature (we decided to use another term to avoid confusion with food webs’ network structure). Food web shape describes the distribution of biomass along the three main food chain compartments: plants, primary and secondary consumers. Food webs can have a pyramid or cascade shape, meaning that biomass can be either distributed alternatively or not along the food chain and can be bottom, middle (except for pyramids) or top-heavy (BH, MH or TH), which described the position of the compartment with the highest biomass. In other words, if we order a community compartments (P for producers, H for herbivores and C for secondary consumers) according to their total biomass, if the result is P-H-C, the community has a BH pyramidal shape, P-C-H gives a BH cascade shape, H-C-P and H-P-C both represent MH cascade shape, C-H-P is a TH pyramidal shape (also called inverted pyramid of biomass) and C-P-H is a TH cascade-shaped community.

## 3 Results

A total of 4589 food webs persisted with all three compartments (plants, primary and secondary consumers). Among these, a vast majority (91.8 %) are bottom heavy (BH), 7.7 % are top heavy (TH) and only 0.5 % are middle heavy (MH). While there are almost as many bottom-heavy cascade-shaped and pyramid-shaped food webs (50.2 % and 49.8 % respectively), top-heavy food webs are predominantly cascade-shaped (94.6 %). The low proportion of TH pyramid-shaped food webs and MH cascade-shaped food webs make the interpretation of these particular results more difficult. We thus focused our interpretation on either only BH food webs and TH cascade-shaped food webs or qualitative comparisons.

The emergence and persistence of the different shapes seem to be related to feedback between food web functioning and structure. While we were not able to see a sizeable difference in structure between cascade-shaped and pyramid-shaped food webs, we do see structural differences between BH, MH and TH food webs (see fig. 2). To better visualize how the different descriptors of structure used in fig. 2 translate in terms of food-web global structure, we represented a typical (median values for connectance and height) food webs for each case (fig. 3), with the average (solid line) and standard deviation (ribbon) for the biomass dynamics of producers (green), herbivores (black), omnivores (light orange) and carnivores (dark orange). Food webs appear in our analysis to be BH by default, yet TH food webs can still persist but they are relatively rare (7.7 %). These TH food webs exhibit greater vertical diversity (i.e., a greater number of trophic levels; see fig. 2 A) allowing them to maintain a higher mean consumer-resource body-mass ratio (see node sizes in the typical food webs, fig. 3). This higher ratio is associated with greater stability (Brose et al. 2006) and large body mass are associated with lower metabolic losses (see model description, appendix 1, section 1 in supplementary material). They are also more complex (higher connectance, see fig. 2 C) and display particular types of interactions. In fact, looking at the motifs profile of food webs, it appears that TH food webs have more omnivory/IGP and exploitative competition motifs (fig. 2 D and G), while BH food webs have on average more apparent competition motifs (fig. 2 D) which seems to be consistent with TH food webs displaying a more complex and functionally diverse secondary consumer compartment. The higher proportion of omnivory is probably related to higher intra-guild predation, allowing for cycling and thus longer biomass residence time in the system.

**Figure 2.**
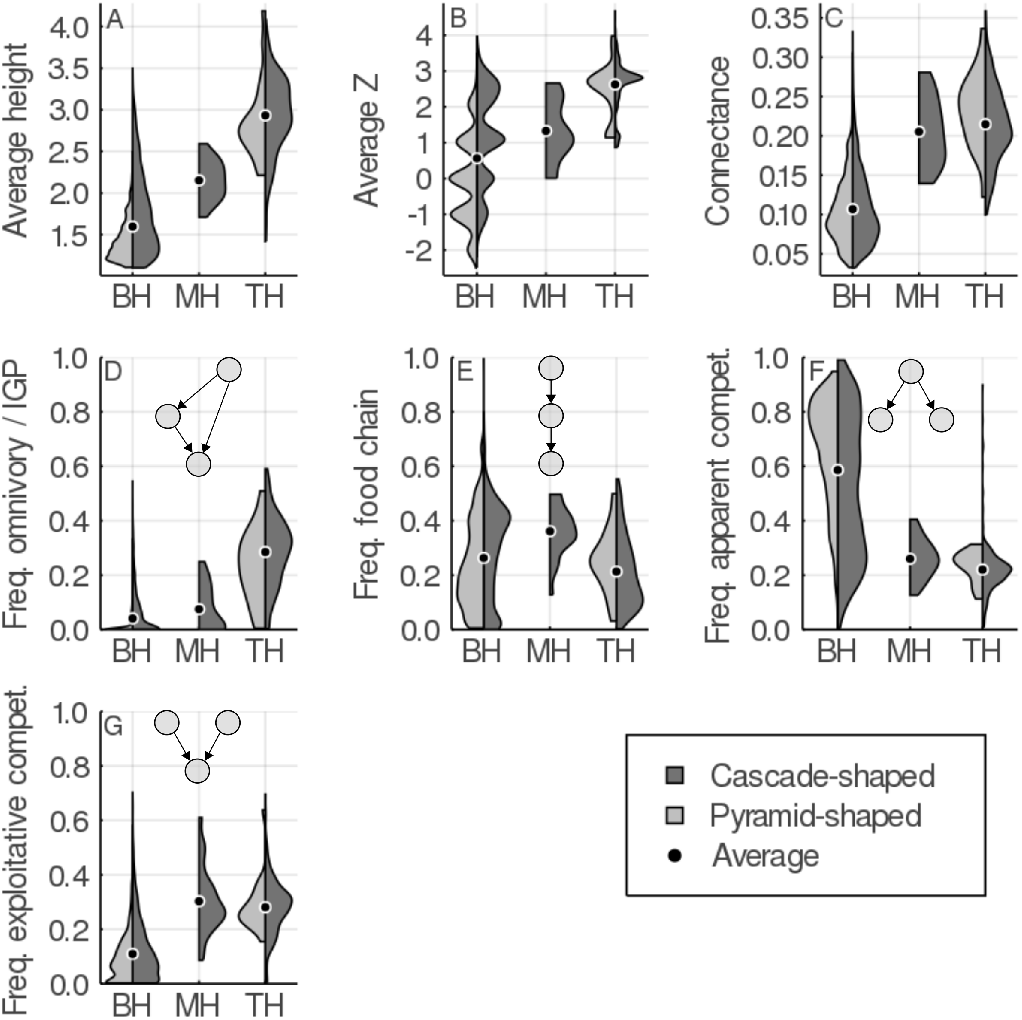
Distribution of some food web structural properties of interest. Here are represented the average food web height (maximum trophic level, A), consumer-resource body-size ratio (log10 scale, B), connectance (C), as well as the average frequency of 4 typical motifs in the food webs. These motifs correspond to typical trophic modules: omnivory (or intra-guild predation, D), linear food chain (E), apparent competition (F) and exploitative competition (G). Motifs are represented on top of these panels.

**Figure 3.**
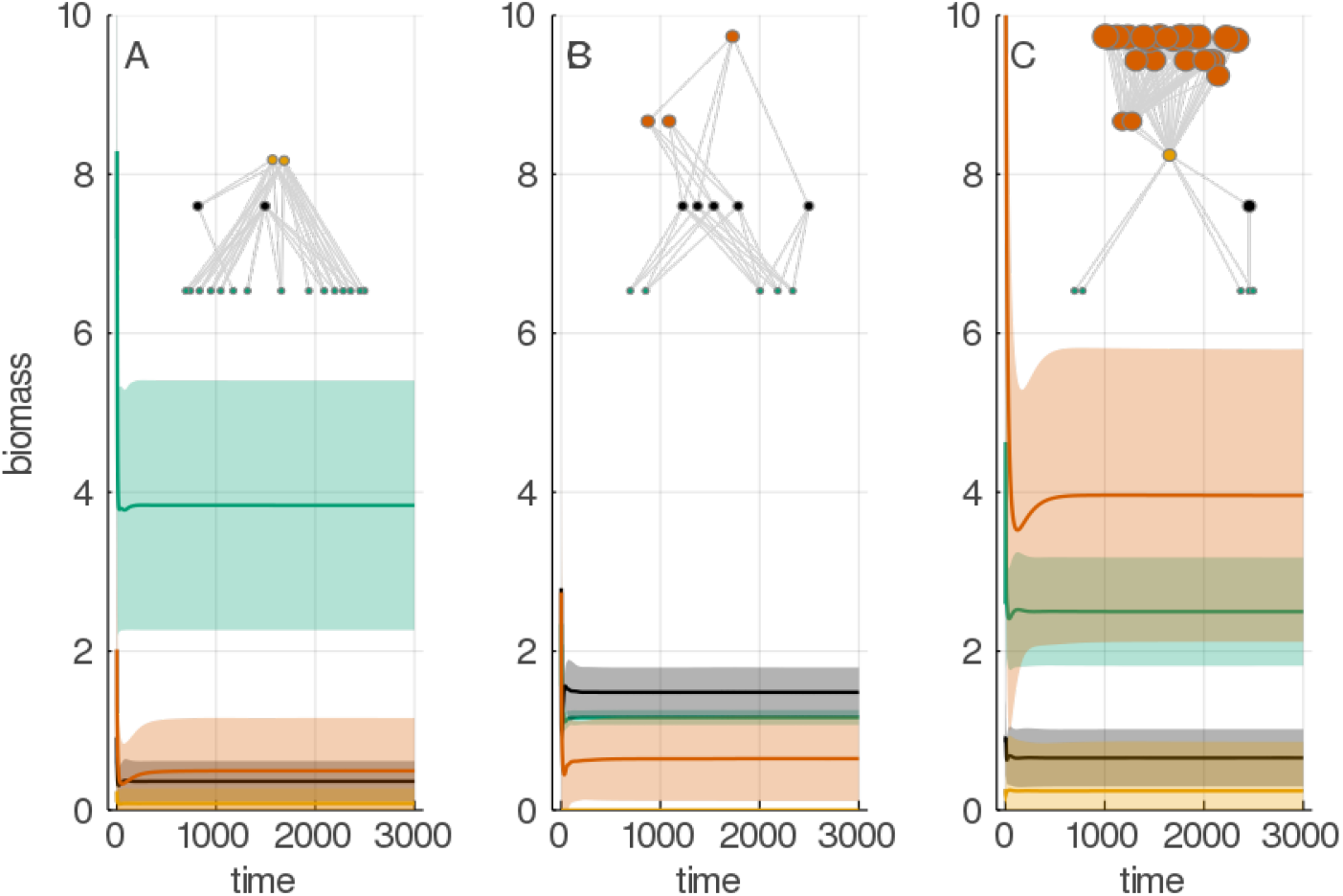
Typical food web structure and associated averaged biomass dynamics. Here are represented the average biomass dynamic for the three main compartments, for all bottom-heavy (A), middle-heavy (B) and top-heavy (C) food webs. For each compartment (plants in green, primary consumers in black and secondary consumers in light or dark orange, respectively for omnivores and carnivores), the total compartment biomass is averaged at each time step (solid line) and are represented with standard deviation (ribbon). A typical food web is plotted above the dynamics, following the same colour code. Nodes sizes represent the body mass of the different species (log10 scaled).

Our results indicate that food webs, comprising both competition and consumption, are in general less efficient at storing the captured biomass than purely competitive communities (4). The dashed line in each panel of figure fig. 4 represents the theoretical biomass to intake regime of community composed of producers only, with no trophic interaction to mediate interspecific competition for the shared nutrients. Intake here represents the food webs total intake, that which is equivalent in our model to primary production. We expect that given energetic constraints (metabolism and imperfect assimilation), communities achieve higher biomass, for the same intake, when there are no trophic interactions involved (*e.g.* a grassland with no consumers). The majority of the food webs meets this expectation, displaying a regime below this reference line. Yet, surprisingly, the producers-only baseline regime can be overshot in some cases (approx. 6.4 % of all food webs). This can happen when food webs total intake is above a certain threshold, the value of this intake threshold being dependent on the shape of the food webs. BH food webs (fig. 4, D and E) or MH (fig. 4, C) have in average a lower biomass to intake regime than TH food webs (fig. 4, A and B), which represent a lower ability to store the captured biomass. For example, for a total intake of 0.75, the theoretical relationship for a grassland gives a biomass of approx. (fig. 4), BH food webs have a biomass largely below this value (< 2.5, fig. 4, D and E) but TH food webs have a biomass between 2.5 and 7 (fig. 4 A and B), higher than BH and in some case than the theoretical baseline. This does not seem to depend on food webs shape (cascade or pyramid), but only on their top heaviness, although BH cascade-shaped food webs seem not to be able to overshoot the baseline even at maximum intake, unlike their cascade-shaped counterpart. Storage efficiency is also strongly correlated to food webs mean body-size consumer-resource ratio (*Z*), at the same intake value, higher *Z* is correlated to higher biomass.

**Figure 4.**
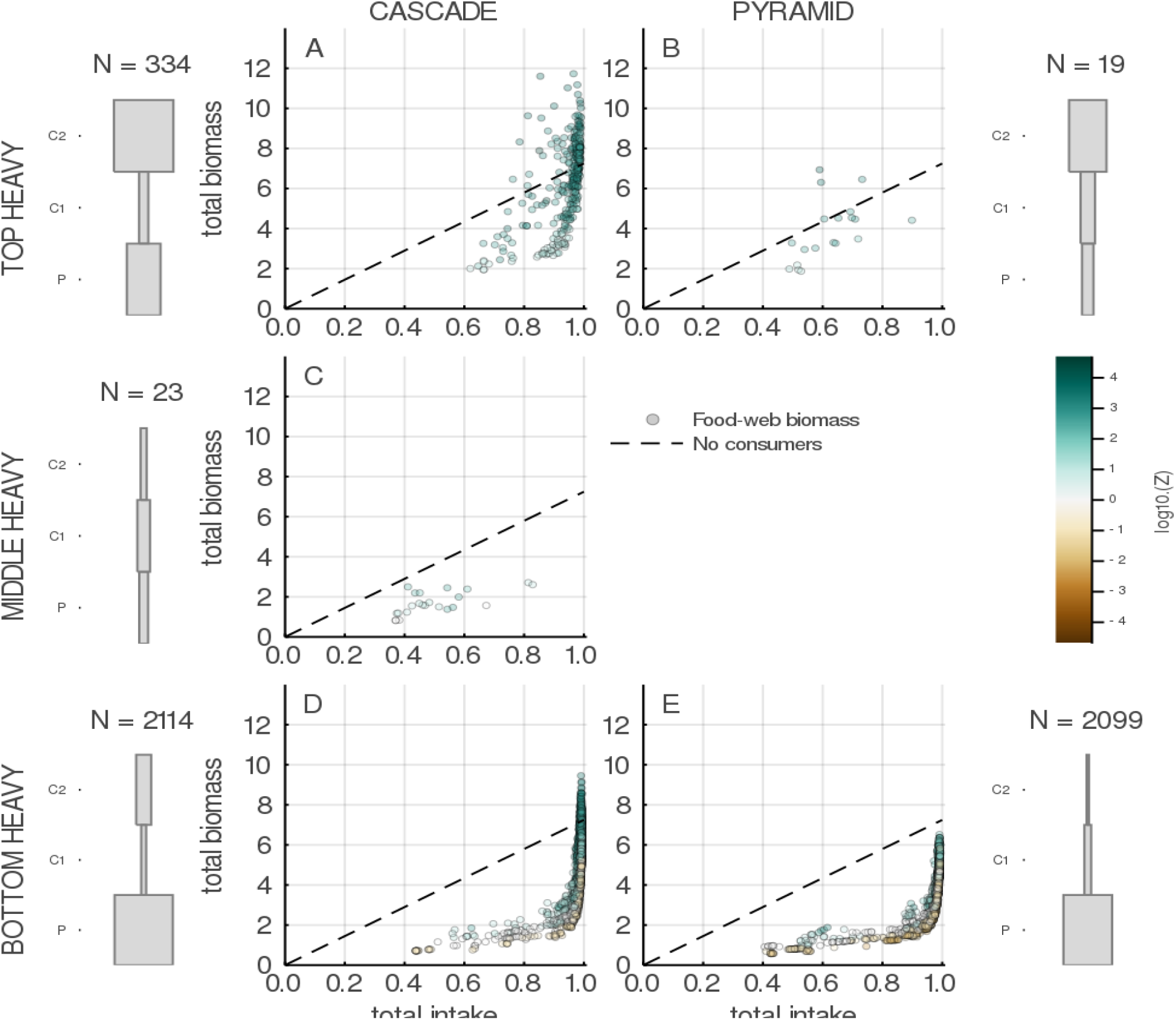
Bottom-heavy food webs have a lower intake-biomass regime on average but can exist at lower intake values and when consumers are on average smaller than their resources (log10(Z) < 0). This figure represents the relationship between food-web total intake and total biomass (each dots represent one food web) for different shapes, namely top-heavy (top, A and B), medium-heavy (middle row, C) and bottom-heavy (bottom, D and E) cascades (left, A, C and D) and pyramids (right, B and E). Dots are coloured according to the decimal logarithm of the average consumer-resource body-size ratio in the food web. The average shape is represented on the left of each plot for cascades and on the right for pyramids along with N, the number of food webs in each panel. The grey boxes represent the mean biomass of the three compartments (Plants P, Herbivores C1 and secondary consumers C2).

Despite the differences mentioned above, food webs appear to all have qualitatively similar diversity-functioning relationships independently of their shape (fig. 6 and fig. 5). Total biomass increases with both animal and plant richness with similar rates for all shapes (fig. 6), although unfortunately there is not enough variation in food webs displaying TH pyramid or MH shapes to analyze their BEF relationship. Looking at the flows within food webs, we see a similar result, with qualitatively analogous relationship for the different shapes, although it appears that TH pyramid-shaped and MH food webs display a lower level of intake (fig. 5). While we reach maximum productivity even for low richness for the BH food webs (fig. 5, bottom row), producers appear more strongly controlled in the TH food webs (fig. 5, top row) – especially cascade-shaped (fig. 5, left column) – and the MH cascades in which herbivores are less regulated (fig. 5, C), resulting in low productivity. Consumption, on the other hand is higher, whatever the level of animal richness in the TH food webs (fig. 5, top row). This higher consumption is driven mainly by higher secondary consumption, in particular higher intra-guild consumption. In TH and BH cascade-shaped food webs, we see that productivity and secondary consumption both increase with richness while primary consumption decreases (fig. 5, A and D). The fact, however, that all richness-functioning relationships are qualitatively similar seems to confirm the existence of a universal diversity effect, albeit more or less strong depending on shape, in food webs.

**Figure 5.**
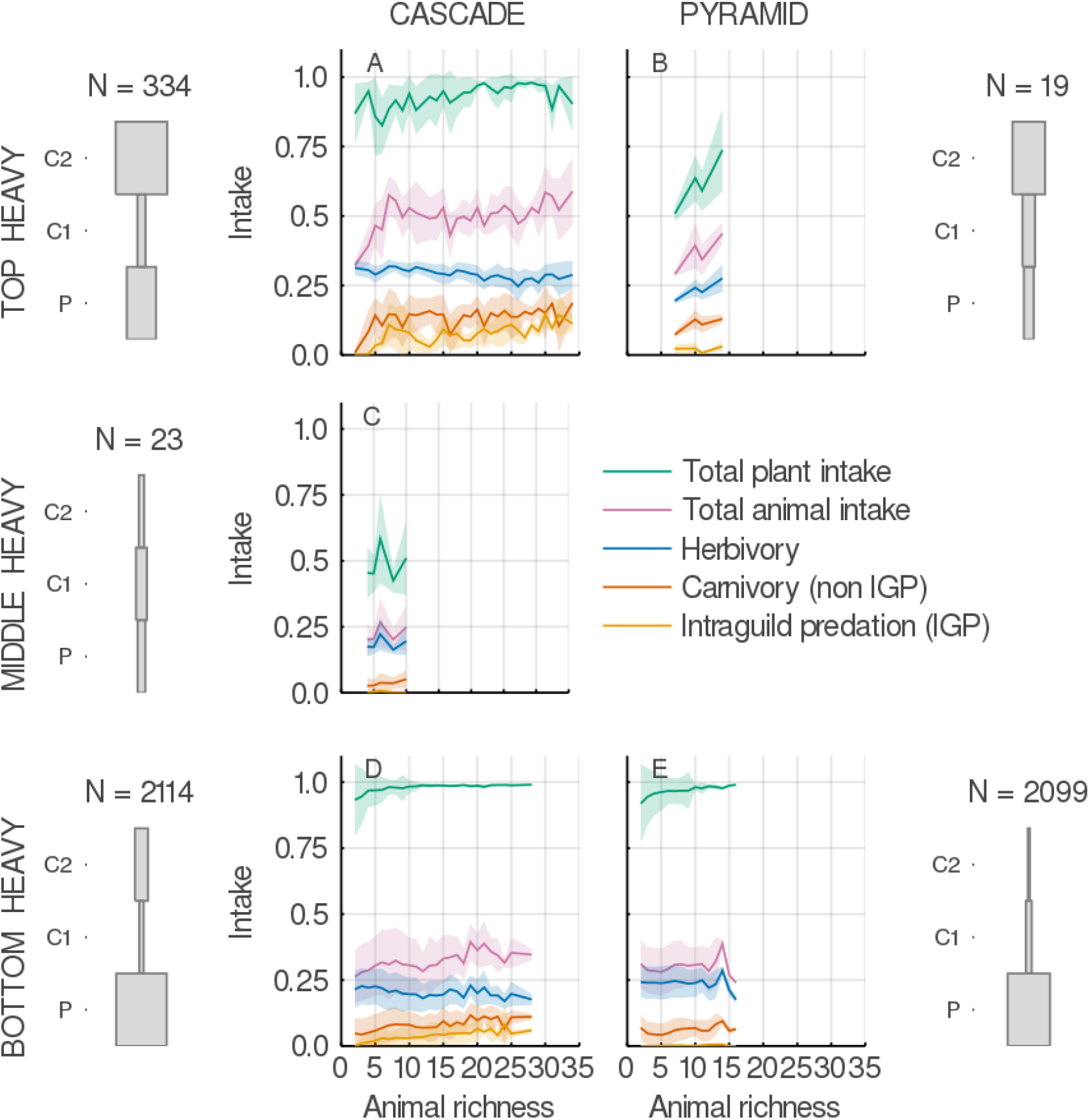
Animal richness - flux relationship in food webs with different shapes. This figures shows the effect of total animal richness on food webs total intake for the different compartments (colour coded, see legend) and the different shapes, namely top-heavy (top, A and B), medium-heavy (middle row, C) and bottom-heavy (bottom, D and E) cascades (left, A, C and D) and pyramids (right, B and E). The solid line represents the average response and the shaded area represent the standard deviation around the mean. The average shape is represented on the left of each plot for cascades and on the right for pyramids along with N, the number of food webs in each panel. The grey boxes represent the mean biomass of the three compartments (Plants P, Herbivores C1 and secondary consumers C2).

**Figure 6.**
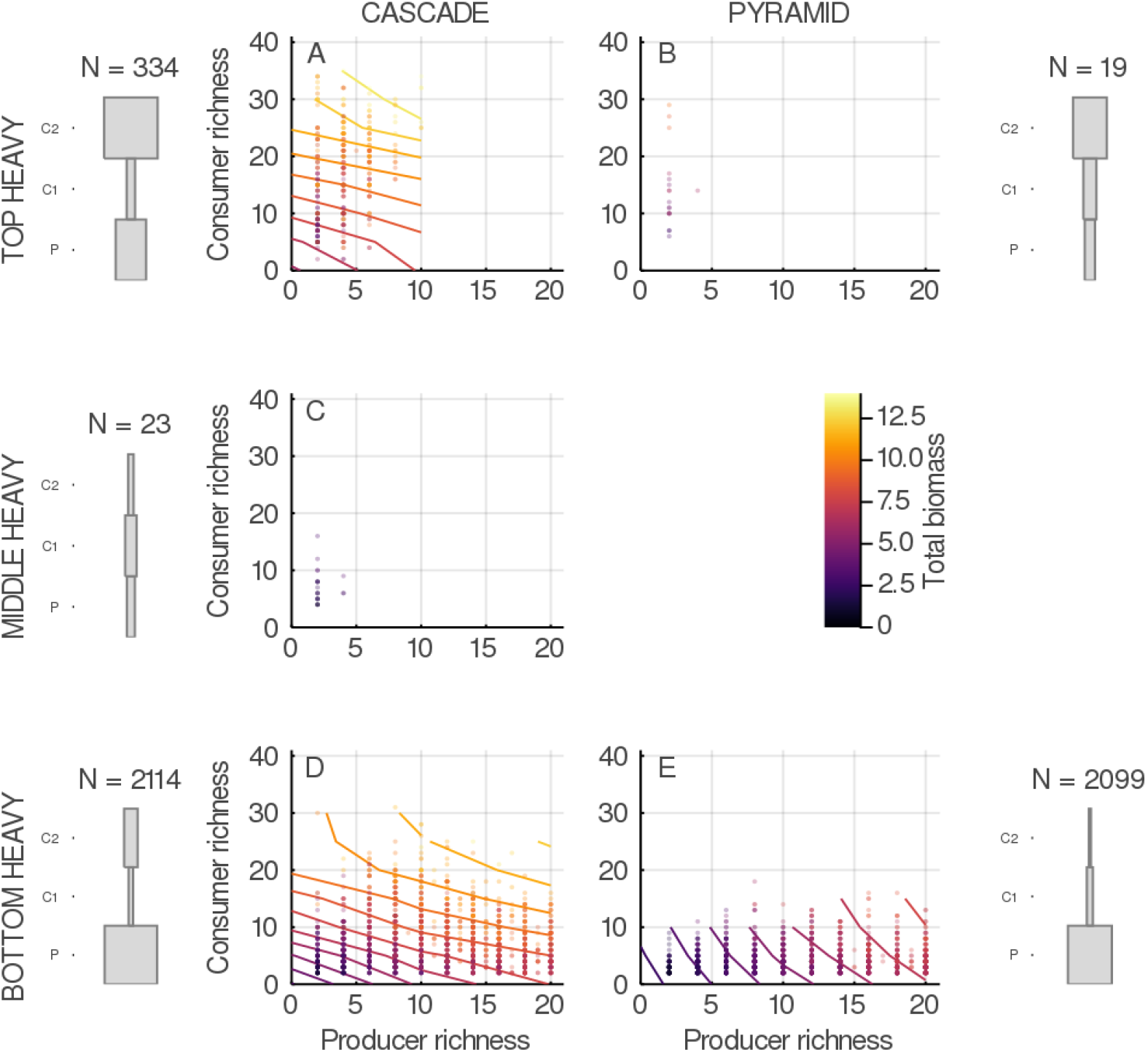
Diversity - total biomass relationships in food webs with different shapes. This figures shows the effect of producers and consumers richness on food webs total biomass for food webs of different shapes, namely top-heavy (top, A and B), medium-heavy (middle row, C) and bottom-heavy (bottom, D and E) cascades (left, A, C and D) and pyramids (right, B and E).

## 4 Discussion

We were able to reproduce predictions typical of different domains of ecology, all of which aim at understanding and predicting ecosystem functioning, but based on different concepts and approaches. On the one hand, the biodiversity-ecosystem functioning (BEF) theory mainly focuses on quantifying the effect of species richness on ecosystem functioning using concepts rooted in competition and predicts a positive and asymptotic BEF relationship (Tilman et al. 1996, 2014; Loreau & Hector 2001). On the other hand, the analysis of biomass transfer in communities represented by trophic chains predicts that communities should be bottom heavy in the absence of biomass or nutrient subsidies (Leroux & Loreau 2008), pyramid-shaped if we adopt a static perspective based on the balance of energy transfers or possibly cascade-shaped if we focus on the dynamics of biomass transfers through trophic interactions (Barbier & Loreau 2019). The methodological framework we used, based on the coupling of the representation of communities through their underlying food webs, and the use of a bioenergetic model to simulate the dynamics of biomass transfers in these trophic networks (Yodzis & Innes 1992; Williams et al. 2007) allows us, firstly, to generate this reconciliation, and secondly to use tools for analyzing food webs structure. The use of these tools is ultimately what allows us to shed new light on the possible link between these different predictions, and thus on the concepts that frame them.

### 4.1 A link between food webs complexity and top heaviness

Through this synthetic framework, we highlight an apparent link between communities’ food web structure and top heaviness. Ecological communities in fact appear to be distributed on a multivariate structural gradient according to their top heaviness. On one side of this gradient, we have a very large majority (91.8 %) of bottom heavy communities, presenting a relatively short trophic chain and a fairly low complexity, in the sense that interactions seem to be only slightly entangled (they display a relatively weaker connectivity and less omnivore/intraguild predation motifs, see fig. 2 and 3). These communities have a high producer species richness at their base, but a low consumer richness (fig. 6), thus displaying consumers feeding low in the food chain and often generalists (which is translated by a high proportion of apparent competition motifs as shown in fig. 2). In other words, these communities are more horizontally than vertically diverse. Moving along this gradient, we see higher and more complex communities at the other end (connectivity is larger and they have a higher proportion of omnivore/intraguild predation motifs, see fig. 2 and 3). Conversely, these communities appear to be richer vertically than horizontally. In the middle, in an intermediate situation, there are very few middle-heavy communities.

When we look at natural ecosystems, we realize that in the same way, they are dominated in most cases by the biomass of heterotrophs (Bar-On et al. 2018). This observation has been at the origin of the bottom-up VS. top-down debate (Wilkinson & Sherratt 2016) as well as much of the work around food chains. In the context of the bioenergetic model, as in nature, energy constraints are such that it is in fact rare to be able to maintain a dominant secondary consumer compartment (Lindeman 1942; McCauley et al. 2018). In addition, the greater risk of extinction faced by predators, especially large top predators (Cardillo 2003; Binzer et al. 2011), add an additional constraint by threatening the persistence of this type of community. This explains why we only find top-heavy food webs at high primary production values (see fig. 4). For this to happen in the absence of external factors such as nutrient supply (Leroux & Loreau 2008), several conditions must be met, that leads to a particularly efficient system in terms of energy transfer (McCauley et al. 2018). This efficiency, translated in our study by a higher capacity to store the biomass produced (see fig. 4), seems to be linked to a particular organization of the interactions, and in turn food web structure, as well as to the allometric scaling of biological rates (fig. 2).

### 4.2 Similar BEF relationships in spite of different efficiencies at storing biomass

This link between food web structure and functioning regime is not surprising if we look at recent results on the analysis of biomass dynamics and the resulting functioning in food webs. The use of a very similar methodological framework revealed a correlation between food webs species richness, height and functioning (Wang & Brose 2017), on the one hand, and between species richness, proportion of intraguild predation and functioning, on the other hand (Wang et al. 2019). The fact that we find these two factors (height and proportion of IGP) correlated with communities requiring a high level of functioning seems therefore quite consistent. What is more surprising is that, while the food web structure does have an effect on the distribution of biomass within the different compartments and on the efficiency in storing the biomass produced, this impact has little effect on the diversity-functioning relationship (see fig. 5 and 6). While we have different vertical vs. horizontal diversity balances — and therefore an equally different competition vs. consumption balance — which should theoretically lead to difference in the BEF relationship (Thébault & Loreau 2006; Duffy et al. 2007) in all cases for which we had enough data to draw conclusions, we have qualitatively very similar (almost identical) diversity-functioning relationships. This is valid whether we look at flows (fig. 5) or total biomass (fig. 5). In the light of the recent results cited above, that focused on the analysis of the diversity-functioning relationship in food webs, this can be explained by a complex interplay between structure, energy efficiency of interactions, biomass distribution, and functioning. The observed effect of intraguild predation in (Wang et al. 2019) or vertical diversity in (Wang & Brose 2017) would only exist in certain ranges of variation in species richness, which can only persist if the community presents a particularly efficient functioning regime, which leads it to be able to store a large biomass in its apical compartment, i.e., in top-heavy form.

### 4.3 Cascade- and pyramid-shaped food webs are only as different as their top heaviness

It is important to note, however, that although our approach yields communities with different shapes (cascade- and pyramid-shaped), communities’ shape does not appear to be related to particular food web structures or to a particular functioning regime (fig. 4). For top-heavy communities, we are potentially limited by the amount of data (only 20 food webs over the total 4589 display an inverted pyramidal shape). The energy constraints are indeed such to maintain a high biomass of carnivores while the biomass of producers is low that it is our understanding that very few communities in the context of our model have been able to persist under these conditions. For bottom-heavy communities, however, we have almost as many pyramid-shaped as we have cascade-shaped communities, and so we do not have this limitation. And there seems to be very little difference between the two, apart from the inability of the bottom-heavy pyramids to overshoot the functioning regime of a purely competitive community. We think this is related to the low relative biomass of carnivores. Indeed, in the cascade-shaped counterpart of the bottom-heavy communities, we note that at high intakes, this baseline is slightly overshot, and that high intakes are generally correlated with high animal diversity. This is what leads us to say that, in the context of our analysis, top heaviness is a more important factor than shape in separating communities according to their functioning regime.

### 4.4 Perspectives

Other factors limit the scope of our results. In our analysis, we took a non-adaptive view of trophic interactions. Faced with the extinction of some or all of its resources, a consumer in our model cannot adapt by switching its diet, which would result in a rewiring of interactions (Petchey et al. 2008; Staniczenko et al. 2010; Gilljam et al. 2015). To disentangle the interplay between rewiring (notably through influence on secondary extinction dynamics) according to different rewiring mechanisms and biomass dynamics seems to us to be a necessary first step for the subsequent inclusion of rewiring in this kind of synthesis. It is possible that by allowing consumers to adapt their diet to the biomass variations of their resource or competitor, we see the biomass dynamics change (Kondoh 2003). This in turn could make it easier to maintain structures that would otherwise not be able to persist given energetic constrains, or to maintain a greater range of diversity for certain structures. This could change, if not qualitatively, at least quantitatively our conclusions. It should also be noted that we took a relatively global approach, and chose not to test the effect that ecosystem type might have on our results. Changing the type of ecosystem, and thus the proportion of vertebrates, ectotherm or endotherm vertebrates (which have different biological rates), could also change the results.

Species face increasing extinction risk, threatening the persistence of ecological processes and functioning, but even in systems where species richness is not affected, the structure of the food web that connects them can be altered by environmental changes (Albouy et al. 2014; Kortsch et al. 2015). Ecological network analyses show us that communities are more than the sum of their parts. To maintain ecosystem stability, functioning and the resulting services to human societies, we need to protect the structure of diversity (McCann 2007; Tylianakis et al. 2010). Preserving ecological network structure requires that we identify, like we did here, the attributes of network structure that are important ecosystem processes, how species contribute to different attributes, how different attributes interacts and how ecosystem processes feedback on structure. This would ease the choice of conservation targets and make conservation more efficient.

We have emphasized the role of food web structure and its importance to understand ecosystem processes. However, accurately sampling ecological networks such as food webs is not an easy task. When sampling food webs, interactions are established using various methods that reflect different ecological realities (Delmas et al. 2019). The ensuing difficulty to evaluate food-web data (Jordano 2016) challenges our understanding of the effect of realistic variations in structure, and in turn our ability to make accurate predictions of functioning based on sampled communities. More work in understanding the mechanisms that underlie the probability of an interaction, to build mechanistic food-web models that produce realistic food webs, and more food-web sampling are still needed. However, we show here that precise information may not necessarily be needed. In fact, in the context of our work, species richness of trophic compartments and the animal to producer ratio can be used to estimate the domain of variation of chain length, motifs distribution, and consumer-resource body-mass ratio, which makes estimating the functioning regime of the community possible.

In conclusion, we show here that food web structure is important in understanding ecosystem functioning, but is also the product of feedbacks between species richness and community functioning. If structure does not appear to be influencing qualitatively the diversity-functioning relationship, it still seems important to other aspects of functioning such as the distribution of biomass along the food chain. This in turn could result in different consequences when facing perturbations, as extinction risk increases with trophic rank. Of course, before understanding the real-world implication, more work is needed. Our analysis lays potentially interesting links, the validity and the generality of the relationships between food web structure, top heaviness and functioning regime should be further tested, and the exact mechanisms underlying the relationships identified.

## Notes

https://osf.io/yuezq/?view_only=2378095a26d9414489fbcd4a72753a27

### Competing Interest Statement

The authors have declared no competing interest.

