## Supplementary information for "Food web structure alters ecological communities top-heaviness with little effect on the biodiversity-functioning relationship"

**2 The allometric diet-breadth model (ADBm) 5**

**3 The Benguela pelagic food web 6**

**4 Breaking down of the diversity-functioning relationship 10**

**References 14**

### 1 THE BIO-ENERGETIC FOOD-WEB MODEL (BEFWM)

#### 1.1 *Biomass dynamics in the BEFW model*

The bio-energetic food web model (Yodzis & Innes 1992; Williams et al. 2007) driving equation is described below (1). The first term describes how producers, at the most basal level of the food web, are responsible for biomass growth (thus making $r_i G_i(N) B_i = 0$ for non-producers species). All producers use the same nutrients ($N$) to produce biomass, sustaining the system. This is described in greater details in the section *Growth through nutrient intake*. Biomass is then transferred through each trophic interaction (second term and third terms of 1) through a multi-species functional response (see section *The multi-species functional response*). Finally, biomass is loss through metabolism (fourth term of 1).

$$B_i' = r_i G_i(N) B_i - \sum_{\text{j = consumers}} \frac{x_j y_j B_j F_{\text{ji}}}{e_{\text{ji}}} + \sum_{\text{k = resources}} x_i y_i B_i F_{\text{ik}} - x_i B_i \quad (1)$$

In 1 and the paragraphs below, $i$ represent the focus species, $j$ represent its consumers, $k$ its resources and $l$ the nutrients. The state variable $B_i$ is the biomass of population $i$. The biomass dependant growth rate of producers is described by the first term of the equation: $r_i G_i(N) B_i$, where $r_i$ is the species intrinsic growth rate and $G_i(N)$ is its net growth rate and depends on nutrients ($N$) concentration (described below in the paragraph Growth through nutrient intake). The second and third terms of the equation are the biomass transfers (respectively loss and gain) through trophic interactions between population $i$, its consumers $j$ and resources $k$. It depends on the consumer ($j$ or $i$) mass-specific metabolic rate ($x$), its maximum consumption rate relative to its metabolic rate ($y$), a multi-resource functional response ($F$, described in more details below in the paragraph *Consumption: the multi-species functional response*) and $i$'s assimilation efficiency when consuming population $j$ ($e_{ij}$). Finally, all species lose biomass through metabolism, this is described by the last term of 1. The values chosen for all parameters are given in 1.

**Table S 1** Values used for the parameters of the bioenergetic model. The metabolic rate $x$ is allometrically scaled and as such is different for each species ($a_x$ is 0.88 for vertebrates and 0.314 for invertebrates, $M_p$ represent the body mass of the smallest producer and $M_i$ the body mass of the focus species).

| Parameter | Symbol | Value |
|---|---|---|
| Intrinsic growth rate | $r_i$ | 1 |
| Metabolic rate | $x_i$ | $a_x(M_i/M_p)^{-0.25}$ |
| Maximum assimilation rate | $y_i$ | 8 for invertebrates, 4 for vertebrates |
| Assimilation efficiency | $e_j i$ | 0.85 for carnivory links, 0.45 otherwise |
| Plants half-saturation densities | $K_{\mathrm{li}}$ | $\mathcal{U} \sim [0.15, 0.20]$ |
| Turnover rate | $D$ | 0.25 |
| Supply | $S_l$ | 4 for each nutrient |
| Nutrient content in plant biomass | $c_l$ | $c_1 = 1$ and $c_2 = 0.5$ |
| Preference | $\omega_{\mathrm{ji}}$ | $1/n$ where $n$ is the number of resources |
| Consumers half-saturation densities | $B_0$ | 0.5 |
| Hill exponent | $h$ | 2 |
| Predator interference | $c$ | 0 |

##### 1.1.1 *Growth through nutrient intake*

The nutrient intake model (Tilman et al. 1996; Brose et al. 2005) describes the relationship between nutrient concentration and producers biomass, and is formalized by the following equation:

$$G_i(N) = MIN(\frac{N_1}{K_{1\mathrm{i}} + N_1}, \frac{N_2}{K_{2\mathrm{i}} + N_2}) \tag{2}$$

In 2 $N_1$ and $N_2$ are the respective concentrations in the environment of the two nutrients shared by all plant species. Competition between plants emerge from this model through the difference in the plants half-saturations densities ($K_{li}$) for the two nutrients. The concentration of the two nutrients is determined by the following equation:

$$N_l' = D(S_l - N_l) - \sum_{i=1}^{n}(c_{li} r_i G_i(N) B_i) \tag{3}$$

where $D$ is the turnover rate relative to the time scale of the system (determined by the intrinsic growth rate), $S_i$ is the supply concentration and $c_{li}$ is the concentration of nutrient $l$ in species $i$, which determines which of the two nutrient species need the most. If plants intrinsic growth rate $r$, metabolic rate $x$ and concentration for each nutrients $c_1$ and $c_2$ are equal, then the hierarchy of competition is primarily determined by species half saturation for $K_{1i}$ (the smaller the stronger competitor $i$ is) if $c_1 > c_2$ and conversely.

##### 1.1.2 *The multi-species functional response*

The second and third terms of 1 describe respectively the amount of biomass lost and gained through consuming or being consumed by other species. While $(x_i y_i B_i)/e_{ji}$ determines the metabolic-dependant efficiency of $i$ at consuming $j$, $F_{ji}$ express the fraction of this efficiency rate that is actually achieved by $j$ when consuming $i$. This fraction, namely the functional response, is expressed through the following equation:

$$F_{ik} = \frac{\omega_{ik} B_k^h}{B_0^h + c B_i B_0^h + \sum_{r = \text{resources}} \omega_{ir} B_r^h} \tag{4}$$

where $\omega_{ik}$ quantify the specialization of $i$ towards $k$, that is the fraction of $i$'s diet (represented by the maximum consumption rate $y_i$) targeted to eating $k$. The parameter $h$, the Hill exponent, controls how the functional response will saturate in response to an increase in $k$'s biomass ($B_k$). Hill coefficient is bound between 1 (Holling type II functional response; Holling 1959) and 2 (Holling type III functional response, Holling 1959; Real 1977), but can take any value in between, making the saturation curve more or less sigmoid. Predator interference can also be implemented in this functional response by making $c > 0$, which expresses the density-dependant control of predator on themselves (DeAngelis et al. 1975). Finally, $B_0$ represent the half-saturation density.

### 2 THE ALLOMETRIC DIET-BREADTH MODEL (ADBM)

To predict interactions between species, the ADBm (Petchey et al. 2008) works in two steps. First it calculates the profitability ($P_{\mathrm{ij}}$, or rate of energy intake) for each pair of species in the community. Then, it selects the links that maximize it. Profitability is expressed as:

$$P_{\mathrm{ij}} = \frac{\sum_{\mathrm{i}=1} \lambda_{\mathrm{ij}} E_i}{1 + \sum_{\mathrm{i}=1} \lambda_{\mathrm{ij}} H_{\mathrm{ij}}} \tag{5}$$

where, $E_i$ is the net energy gained by $j$ when consuming $i$ and scales linearly with $i$'s body size ($M_i$): $E_i = eM_i$. Profitability also depends on the encounter rates $\lambda_{\mathrm{ij}}$ ($\lambda_{\mathrm{ij}} = N_i * A_{ij}$) which depends on the density of the resource species ($N_i = nM_i$) and the attack rate ($A_{ij} = aM_i^{a_i} M_j^{a_j}$). Finally, profitability is also influenced by the handling time $H_{\mathrm{ij}}$. We chose to implement the "ratio" method for estimating handling time as it is supposed to yield more accurate results. In this formulation, handling time is estimated differently depending on the body-size ratio between a consumer and its potential prey. If the size different is too big (bigger than a chosen threshold $b$) then we assume that $i$ is not able to consume $j$. This is expressed by having $H_{ij} = h/(b - (M_i/M_j))$ if $M_i/M_j > b$ and $H_{ij} = \inf$ otherwise. Parameters values are presented below in the 2.

**Table S 2** Values used for the parameters of the allometric diet breadth model. For more details and references for the parameters values used, see Petchey et al., 2008.

| Parameter | Symbol | Value |
|---|---|---|
| Allometric constant for attack rate | $a$ | 0.0189 |
| Consumers allometric coefficient for attack rate | $a_i$ | −0.491 |
| Resources allometric coefficient for attack rate | $a_j$ | −0.465 |
| Allometric constant for handling time | $h$ | 1 |
| Threshold for handling time | $b$ | 0.401 |

### 3 THE BENGUELA PELAGIC FOOD WEB

We used body mass data from species of the Benguela upwelling system, as described in Yodzis (1998) and Brose et al. (2016). From the complete list of interactions, we only kept herbivorous and predacious links and discarded bacterivorous links. Heterotrophic producers were identified as the resource of herbivorous interactions. We sampled body masses from species mean body mass data, and recorded the metabolic type associated with it. The following table provides a list of all species with their body mass and metabolic type.

**Table S 3** Herbivorous and predacious interactions in the Benguela pelagic food web associated with species common names, body masses and metabolic classes.

| Interaction type | Consumer species | Consumer metab | Consumer mass | Resource species | Resource metab | Resource mass |
|---|---|---|---|---|---|---|
| herbivorous | Bacteria | heterotrophic bacteria | 1.0e-8 | Phytoplankton | photo-autotroph | 0.0001 |
| predacious | Benthic carnivores | invertebrate | 10 | Benthic filter feeders | invertebrate | 10 |
| herbivorous | Microzooplankton | invertebrate | 0.0001 | Phytoplankton | photo-autotroph | 0.0001 |
| predacious | Microzooplankton | invertebrate | 0.0001 | Microzooplankton | invertebrate | 0.0001 |
| herbivorous | Mesozooplankton | invertebrate | 0.01 | Phytoplankton | photo-autotroph | 0.0001 |
| predacious | Mesozooplankton | invertebrate | 0.01 | Microzooplankton | invertebrate | 0.0001 |
| herbivorous | Macrozooplankton | invertebrate | 1 | Phytoplankton | photo-autotroph | 0.0001 |
| predacious | Macrozooplankton | invertebrate | 1 | Mesozooplankton | invertebrate | 0.01 |
| predacious | Macrozooplankton | invertebrate | 1 | Macrozooplankton | invertebrate | 1 |
| herbivorous | Gelatinous zooplankton | invertebrate | 100 | Phytoplankton | photo-autotroph | 0.0001 |
| predacious | Gelatinous zooplankton | invertebrate | 100 | Microzooplankton | invertebrate | 0.0001 |
| predacious | Gelatinous zooplankton | invertebrate | 100 | Macrozooplankton | invertebrate | 1 |
| herbivorous | Anchovy | ectotherm vertebrate | 11.5 | Phytoplankton | photo-autotroph | 0.0001 |
| predacious | Anchovy | ectotherm vertebrate | 11.5 | Microzooplankton | invertebrate | 0.0001 |
| predacious | Anchovy | ectotherm vertebrate | 11.5 | Macrozooplankton | invertebrate | 1 |
| herbivorous | Pilchard | ectotherm vertebrate | 280 | Phytoplankton | photo-autotroph | 0.0001 |
| predacious | Pilchard | ectotherm vertebrate | 280 | Macrozooplankton | invertebrate | 1 |
| predacious | Round herring | ectotherm vertebrate | 215.2 | Macrozooplankton | invertebrate | 1 |
| predacious | Lightfish | ectotherm vertebrate | 4.8 | Macrozooplankton | invertebrate | 1 |
| predacious | Lanternfish | ectotherm vertebrate | 6.9 | Macrozooplankton | invertebrate | 1 |
| herbivorous | Goby | ectotherm vertebrate | 18.6 | Phytoplankton | photo-autotroph | 0.0001 |
| predacious | Goby | ectotherm vertebrate | 18.6 | Macrozooplankton | invertebrate | 1 |
| predacious | Other pelagics | ectotherm vertebrate | 2554.85 | Benthic carnvores | invertebrate | 10 |
| predacious | Other pelagics | ectotherm vertebrate | 2554.85 | Macrozooplankton | invertebrate | 1 |
| predacious | Other pelagics | ectotherm vertebrate | 2554.85 | Gelatinous zooplankton | invertebrate | 100 |
| predacious | Horse mackerel | ectotherm vertebrate | 5104.9 | Lanternfish | ectotherm vertebrate | 6.9 |
| predacious | Horse mackerel | ectotherm vertebrate | 5104.9 | Benthic carnivores | invertebrate | 10 |
| predacious | Horse mackerel | ectotherm vertebrate | 5104.9 | Macrozooplankton | invertebrate | 1 |
| predacious | Chub mackerel | ectotherm vertebrate | 3259.5 | Lanternfish | ectotherm vertebrate | 6.9 |
| predacious | Chub mackerel | ectotherm vertebrate | 3259.5 | Benthic carnivores | invertebrate | 10 |
| predacious | Chub mackerel | ectotherm vertebrate | 3259.5 | Macrozooplankton | invertebrate | 1 |
| predacious | Chub mackerel | ectotherm vertebrate | 3259.5 | Round herring | ectotherm vertebrate | 215.2 |
| predacious | Other groundfish | ectotherm vertebrate | 13127 | Round herring | ectotherm vertebrate | 215.2 |

| | | | | | | |
|---|---|---|---|---|---|---|
| predacious | Other groundfish | ectotherm vertebrate | 13127 | Lightfish | ectotherm vertebrate | 4.8 |
| predacious | Other groundfish | ectotherm vertebrate | 13127 | Lanternfish | ectotherm vertebrate | 6.9 |
| predacious | Other groundfish | ectotherm vertebrate | 13127 | Goby | ectotherm vertebrate | 18.6 |
| predacious | Other groundfish | ectotherm vertebrate | 13127 | Other pelagics | ectotherm vertebrate | 2554.85 |
| predacious | Other groundfish | ectotherm vertebrate | 13127 | Other groundfish | ectotherm vertebrate | 13127 |
| predacious | Other groundfish | ectotherm vertebrate | 13127 | Hakes | ectotherm vertebrate | 22994.5 |
| predacious | Other groundfish | ectotherm vertebrate | 13127 | Squid | invertebrate | 40 |
| predacious | Other groundfish | ectotherm vertebrate | 13127 | Benthic carnivores | invertebrate | 10 |
| predacious | Other groundfish | ectotherm vertebrate | 13127 | Mesozooplankton | invertebrate | 0.01 |
| predacious | Other groundfish | ectotherm vertebrate | 13127 | Macrozooplankton | invertebrate | 1 |
| predacious | Other groundfish | ectotherm vertebrate | 13127 | Anchovy | ectotherm vertebrate | 11.5 |
| predacious | Hakes | ectotherm vertebrate | 22994.5 | Pilchard | ectotherm vertebrate | 280 |
| predacious | Hakes | ectotherm vertebrate | 22994.5 | Round herring | ectotherm vertebrate | 215.2 |
| predacious | Hakes | ectotherm vertebrate | 22994.5 | Lightfish | ectotherm vertebrate | 4.8 |
| predacious | Hakes | ectotherm vertebrate | 22994.5 | Lanternfish | ectotherm vertebrate | 6.9 |
| predacious | Hakes | ectotherm vertebrate | 22994.5 | Goby | ectotherm vertebrate | 18.6 |
| predacious | Hakes | ectotherm vertebrate | 22994.5 | Horse mackerel | ectotherm vertebrate | 5104.9 |
| predacious | Hakes | ectotherm vertebrate | 22994.5 | Chub mackerel | ectotherm vertebrate | 3259.5 |
| predacious | Hakes | ectotherm vertebrate | 22994.5 | Other groundfish | ectotherm vertebrate | 13127 |
| predacious | Hakes | ectotherm vertebrate | 22994.5 | Hakes | ectotherm vertebrate | 22994.5 |
| predacious | Hakes | ectotherm vertebrate | 22994.5 | Squid | invertebrate | 40 |
| predacious | Hakes | ectotherm vertebrate | 22994.5 | Mesozooplankton | invertebrate | 0.01 |
| predacious | Hakes | ectotherm vertebrate | 22994.5 | Macrozooplankton | invertebrate | 1 |
| predacious | Hakes | ectotherm vertebrate | 22994.5 | Anchovy | ectotherm vertebrate | 11.5 |
| predacious | Squid | invertebrate | 40 | Pilchard | ectotherm vertebrate | 280 |
| predacious | Squid | invertebrate | 40 | Round herring | ectotherm vertebrate | 215.2 |
| predacious | Squid | invertebrate | 40 | Lightfish | ectotherm vertebrate | 4.8 |
| predacious | Squid | invertebrate | 40 | Goby | ectotherm vertebrate | 18.6 |
| predacious | Squid | invertebrate | 40 | Horse mackerel | ectotherm vertebrate | 5104.9 |
| predacious | Squid | invertebrate | 40 | Other groundfish | ectotherm vertebrate | 13127 |
| predacious | Squid | invertebrate | 40 | Hakes | ectotherm vertebrate | 22994.5 |
| predacious | Squid | invertebrate | 40 | Squid | invertebrate | 40 |
| predacious | Squid | invertebrate | 40 | Benthic carnivores | invertebrate | 10 |
| predacious | Squid | invertebrate | 40 | Macrozooplankton | invertebrate | 1 |
| predacious | Squid | invertebrate | 40 | Anchovy | ectotherm vertebrate | 11.5 |
| predacious | Tunas | ectotherm vertebrate | 909000 | Pilchard | ectotherm vertebrate | 280 |
| predacious | Tunas | ectotherm vertebrate | 909000 | Round herring | ectotherm vertebrate | 215.2 |
| predacious | Tunas | ectotherm vertebrate | 909000 | Lightfish | ectotherm vertebrate | 4.8 |
| predacious | Tunas | ectotherm vertebrate | 909000 | Lanternfish | ectotherm vertebrate | 6.9 |
| predacious | Tunas | ectotherm vertebrate | 909000 | Goby | ectotherm vertebrate | 18.6 |
| predacious | Tunas | ectotherm vertebrate | 909000 | Other pelagics | ectotherm vertebrate | 2554.85 |
| predacious | Tunas | ectotherm vertebrate | 909000 | Horse mackerel | ectotherm vertebrate | 5104.9 |
| predacious | Tunas | ectotherm vertebrate | 909000 | Chub mackerel | ectotherm vertebrate | 3259.5 |
| predacious | Tunas | ectotherm vertebrate | 909000 | Hakes | ectotherm vertebrate | 22994.5 |
| predacious | Tunas | ectotherm vertebrate | 909000 | Squid | invertebrate | 40 |
| predacious | Tunas | ectotherm vertebrate | 909000 | Benthic carnivores | invertebrate | 10 |
| predacious | Tunas | ectotherm vertebrate | 909000 | Anchovy | ectotherm vertebrate | 11.5 |
| predacious | Snoek | ectotherm vertebrate | 13012.1 | Pilchard | ectotherm vertebrate | 280 |
| predacious | Snoek | ectotherm vertebrate | 13012.1 | Round herring | ectotherm vertebrate | 215.2 |
| predacious | Snoek | ectotherm vertebrate | 13012.1 | Lightfish | ectotherm vertebrate | 4.8 |
| predacious | Snoek | ectotherm vertebrate | 13012.1 | Lanternfish | ectotherm vertebrate | 6.9 |
| predacious | Snoek | ectotherm vertebrate | 13012.1 | Goby | ectotherm vertebrate | 18.6 |
| predacious | Snoek | ectotherm vertebrate | 13012.1 | Horse mackerel | ectotherm vertebrate | 5104.9 |
| predacious | Snoek | ectotherm vertebrate | 13012.1 | Hakes | ectotherm vertebrate | 22994.5 |
| predacious | Snoek | ectotherm vertebrate | 13012.1 | Squid | invertebrate | 40 |
| predacious | Snoek | ectotherm vertebrate | 13012.1 | Benthic carnivores | invertebrate | 10 |

| | | | | | | |
|---|---|---|---|---|---|---|
| predacious | Snoek | ectotherm vertebrate | 13012.1 | Macrozooplankton | invertebrate | 1 |
| predacious | Snoek | ectotherm vertebrate | 13012.1 | Anchovy | ectotherm vertebrate | 11.5 |
| predacious | Kob | ectotherm vertebrate | 68000 | Pilchard | ectotherm vertebrate | 280 |
| predacious | Kob | ectotherm vertebrate | 68000 | Goby | ectotherm vertebrate | 18.6 |
| predacious | Kob | ectotherm vertebrate | 68000 | Horse mackerel | ectotherm vertebrate | 5104.9 |
| predacious | Kob | ectotherm vertebrate | 68000 | Chub mackerel | ectotherm vertebrate | 3259.5 |
| predacious | Kob | ectotherm vertebrate | 68000 | Other groundfish | ectotherm vertebrate | 13127 |
| predacious | Kob | ectotherm vertebrate | 68000 | Hakes | ectotherm vertebrate | 22994.5 |
| predacious | Kob | ectotherm vertebrate | 68000 | Squid | invertebrate | 40 |
| predacious | Kob | ectotherm vertebrate | 68000 | Kob | ectotherm vertebrate | 68000 |
| predacious | Kob | ectotherm vertebrate | 68000 | Benthic carnivores | invertebrate | 10 |
| predacious | Kob | ectotherm vertebrate | 68000 | Macrozooplankton | invertebrate | 1 |
| predacious | Kob | ectotherm vertebrate | 68000 | Anchovy | ectotherm vertebrate | 11.5 |
| predacious | Yellowtail | ectotherm vertebrate | 82040.3 | Pilchard | ectotherm vertebrate | 280 |
| predacious | Yellowtail | ectotherm vertebrate | 82040.3 | Round herring | ectotherm vertebrate | 215.2 |
| predacious | Yellowtail | ectotherm vertebrate | 82040.3 | Goby | ectotherm vertebrate | 18.6 |
| predacious | Yellowtail | ectotherm vertebrate | 82040.3 | Other pelagics | ectotherm vertebrate | 2554.85 |
| predacious | Yellowtail | ectotherm vertebrate | 82040.3 | Horse mackerel | ectotherm vertebrate | 5104.9 |
| predacious | Yellowtail | ectotherm vertebrate | 82040.3 | Chub mackerel | ectotherm vertebrate | 3259.5 |
| predacious | Yellowtail | ectotherm vertebrate | 82040.3 | Other groundfish | ectotherm vertebrate | 13127 |
| predacious | Yellowtail | ectotherm vertebrate | 82040.3 | Squid | invertebrate | 40 |
| predacious | Yellowtail | ectotherm vertebrate | 82040.3 | Macrozooplankton | invertebrate | 1 |
| predacious | Yellowtail | ectotherm vertebrate | 82040.3 | Anchovy | ectotherm vertebrate | 11.5 |
| predacious | Geelbek | ectotherm vertebrate | 26127.38 | Pilchard | ectotherm vertebrate | 280 |
| predacious | Geelbek | ectotherm vertebrate | 26127.38 | Round herring | ectotherm vertebrate | 215.2 |
| predacious | Geelbek | ectotherm vertebrate | 26127.38 | Goby | ectotherm vertebrate | 18.6 |
| predacious | Geelbek | ectotherm vertebrate | 26127.38 | Other pelagics | ectotherm vertebrate | 2554.85 |
| predacious | Geelbek | ectotherm vertebrate | 26127.38 | Horse mackerel | ectotherm vertebrate | 5104.9 |
| predacious | Geelbek | ectotherm vertebrate | 26127.38 | Other groundfish | ectotherm vertebrate | 13127 |
| predacious | Geelbek | ectotherm vertebrate | 26127.38 | Hakes | ectotherm vertebrate | 22994.5 |
| predacious | Geelbek | ectotherm vertebrate | 26127.38 | Squid | invertebrate | 40 |
| predacious | Geelbek | ectotherm vertebrate | 26127.38 | Benthic carnivores | invertebrate | 10 |
| predacious | Geelbek | ectotherm vertebrate | 26127.38 | Anchovy | ectotherm vertebrate | 11.5 |
| predacious | Whales and Dolphins | endotherm vertebrate | 82000 | Pilchard | ectotherm vertebrate | 280 |
| predacious | Whales and Dolphins | endotherm vertebrate | 82000 | Round herring | ectotherm vertebrate | 215.2 |
| predacious | Whales and Dolphins | endotherm vertebrate | 82000 | Lanternfish | ectotherm vertebrate | 6.9 |
| predacious | Whales and Dolphins | endotherm vertebrate | 82000 | Goby | ectotherm vertebrate | 18.6 |
| predacious | Whales and Dolphins | endotherm vertebrate | 82000 | Other pelagics | ectotherm vertebrate | 2554.85 |
| predacious | Whales and Dolphins | endotherm vertebrate | 82000 | Horse mackerel | ectotherm vertebrate | 5104.9 |
| predacious | Whales and Dolphins | endotherm vertebrate | 82000 | Hakes | ectotherm vertebrate | 22994.5 |
| predacious | Whales and Dolphins | endotherm vertebrate | 82000 | Squid | invertebrate | 40 |
| predacious | Whales and Dolphins | endotherm vertebrate | 82000 | Macrozooplankton | invertebrate | 1 |
| predacious | Whales and Dolphins | endotherm vertebrate | 82000 | Anchovy | ectotherm vertebrate | 11.5 |
| predacious | Birds | endotherm vertebrate | 2287 | Pilchard | ectotherm vertebrate | 280 |
| predacious | Birds | endotherm vertebrate | 2287 | Round herring | ectotherm vertebrate | 215.2 |
| predacious | Birds | endotherm vertebrate | 2287 | Lightfish | ectotherm vertebrate | 4.8 |
| predacious | Birds | endotherm vertebrate | 2287 | Lanternfish | ectotherm vertebrate | 6.9 |
| predacious | Birds | endotherm vertebrate | 2287 | Goby | ectotherm vertebrate | 18.6 |
| predacious | Birds | endotherm vertebrate | 2287 | Other pelagics | ectotherm vertebrate | 2554.85 |
| predacious | Birds | endotherm vertebrate | 2287 | Horse mackerel | ectotherm vertebrate | 5104.9 |
| predacious | Birds | endotherm vertebrate | 2287 | Chub mackerel | ectotherm vertebrate | 3259.5 |
| predacious | Birds | endotherm vertebrate | 2287 | Other groundfish | ectotherm vertebrate | 13127 |
| predacious | Birds | endotherm vertebrate | 2287 | Hakes | ectotherm vertebrate | 22994.5 |
| predacious | Birds | endotherm vertebrate | 2287 | Squid | invertebrate | 40 |
| predacious | Birds | endotherm vertebrate | 2287 | Snoek | ectotherm vertebrate | 13012.1 |
| predacious | Birds | endotherm vertebrate | 2287 | Birds | endotherm vertebrate | 2287 |

| | | | | | | |
|---|---|---|---|---|---|---|
| predacious | Birds | endotherm vertebrate | 2287 | Seals | endotherm vertebrate | 136000 |
| predacious | Birds | endotherm vertebrate | 2287 | Benthic carnivores | invertebrate | 10 |
| predacious | Birds | endotherm vertebrate | 2287 | Mesozooplankton | invertebrate | 0.01 |
| predacious | Birds | endotherm vertebrate | 2287 | Macrozooplankton | invertebrate | 1 |
| predacious | Birds | endotherm vertebrate | 2287 | Anchovy | ectotherm vertebrate | 11.5 |
| predacious | Seals | endotherm vertebrate | 136000 | Pilchard | ectotherm vertebrate | 280 |
| predacious | Seals | endotherm vertebrate | 136000 | Round herring | ectotherm vertebrate | 215.2 |
| predacious | Seals | endotherm vertebrate | 136000 | Lightfish | ectotherm vertebrate | 4.8 |
| predacious | Seals | endotherm vertebrate | 136000 | Lanternfish | ectotherm vertebrate | 6.9 |
| predacious | Seals | endotherm vertebrate | 136000 | Goby | ectotherm vertebrate | 18.6 |
| predacious | Seals | endotherm vertebrate | 136000 | Other pelagics | ectotherm vertebrate | 2554.85 |
| predacious | Seals | endotherm vertebrate | 136000 | Horse mackerel | ectotherm vertebrate | 5104.9 |
| predacious | Seals | endotherm vertebrate | 136000 | Chub mackerel | ectotherm vertebrate | 3259.5 |
| predacious | Seals | endotherm vertebrate | 136000 | Other groundfish | ectotherm vertebrate | 13127 |
| predacious | Seals | endotherm vertebrate | 136000 | Hakes | ectotherm vertebrate | 22994.5 |
| predacious | Seals | endotherm vertebrate | 136000 | Squid | invertebrate | 40 |
| predacious | Seals | endotherm vertebrate | 136000 | Snoek | ectotherm vertebrate | 13012.1 |
| predacious | Seals | endotherm vertebrate | 136000 | Birds | endotherm vertebrate | 2287 |
| predacious | Seals | endotherm vertebrate | 136000 | Sharks | ectotherm vertebrate | 1500 |
| predacious | Seals | endotherm vertebrate | 136000 | Benthic carnivores | invertebrate | 10 |
| predacious | Seals | endotherm vertebrate | 136000 | Anchovy | ectotherm vertebrate | 11.5 |
| predacious | Sharks | ectotherm vertebrate | 1500 | Pilchard | ectotherm vertebrate | 280 |
| predacious | Sharks | ectotherm vertebrate | 1500 | Round herring | ectotherm vertebrate | 215.2 |
| predacious | Sharks | ectotherm vertebrate | 1500 | Lightfish | ectotherm vertebrate | 4.8 |
| predacious | Sharks | ectotherm vertebrate | 1500 | Goby | ectotherm vertebrate | 18.6 |
| predacious | Sharks | ectotherm vertebrate | 1500 | Other pelagics | ectotherm vertebrate | 2554.85 |
| predacious | Sharks | ectotherm vertebrate | 1500 | Horse mackerel | ectotherm vertebrate | 5104.9 |
| predacious | Sharks | ectotherm vertebrate | 1500 | Chub mackerel | ectotherm vertebrate | 3259.5 |
| predacious | Sharks | ectotherm vertebrate | 1500 | Other groundfish | ectotherm vertebrate | 13127 |
| predacious | Sharks | ectotherm vertebrate | 1500 | Hakes | ectotherm vertebrate | 22994.5 |
| predacious | Sharks | ectotherm vertebrate | 1500 | Squid | invertebrate | 40 |
| predacious | Sharks | ectotherm vertebrate | 1500 | Tunas | ectotherm vertebrate | 909000 |
| predacious | Sharks | ectotherm vertebrate | 1500 | Snoek | ectotherm vertebrate | 13012.1 |
| predacious | Sharks | ectotherm vertebrate | 1500 | Kob | ectotherm vertebrate | 68000 |
| predacious | Sharks | ectotherm vertebrate | 1500 | Yellowtail | ectotherm vertebrate | 82040.3 |
| predacious | Sharks | ectotherm vertebrate | 1500 | Geelbek | ectotherm vertebrate | 26127.38 |
| predacious | Sharks | ectotherm vertebrate | 1500 | Whales and Dolphins | endotherm vertebrate | 82000 |
| predacious | Sharks | ectotherm vertebrate | 1500 | Birds | endotherm vertebrate | 2287 |
| predacious | Sharks | ectotherm vertebrate | 1500 | Seals | endotherm vertebrate | 136000 |
| predacious | Sharks | ectotherm vertebrate | 1500 | Sharks | ectotherm vertebrate | 1500 |
| predacious | Sharks | ectotherm vertebrate | 1500 | Benthic carnivores | invertebrate | 10 |
| predacious | Sharks | ectotherm vertebrate | 1500 | Mesozooplankton | invertebrate | 0.01 |
| predacious | Sharks | ectotherm vertebrate | 1500 | Macrozooplankton | invertebrate | 1 |
| predacious | Sharks | ectotherm vertebrate | 1500 | Anchovy | ectotherm vertebrate | 11.5 |

### 4 BREAKING DOWN OF THE DIVERSITY-FUNCTIONING RELATIONSHIP

**Figure S 1** Animal to producer biomass ratio for different levels of diversity. This figures shows the relationship between producers and consumers richness on food webs animal to producer biomass ratio for food webs of different shapes, namely top-heavy (top, A and B), medium-heavy (middle row, C) and bottom-heavy (bottom, D and E) cascades (left, A, C and D) and pyramids (right, B and E).

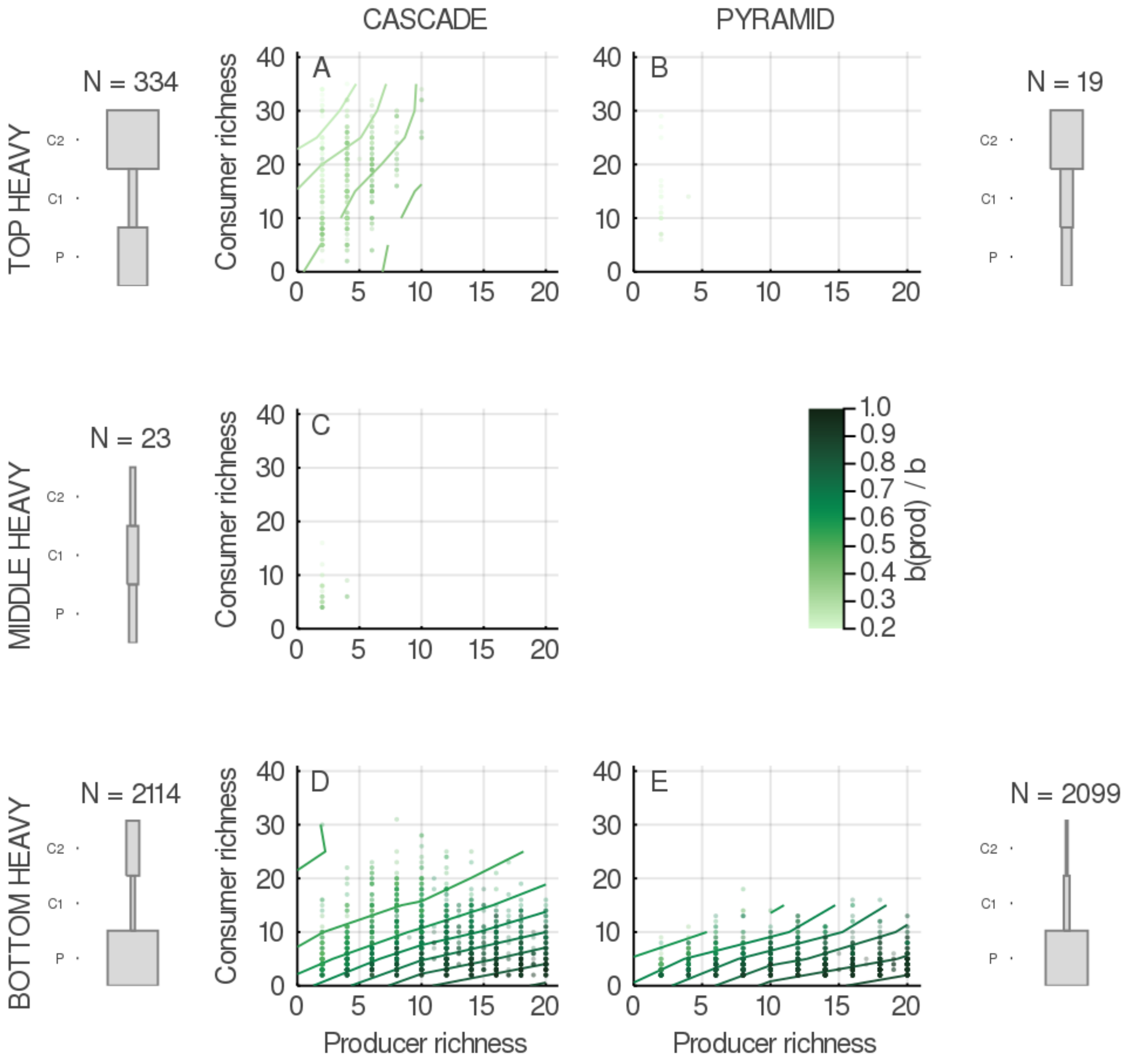

**Figure S 2** Diversity - total primary production relationships in food webs. This figures shows the effect of producers and consumers richness on food webs total primary production for food webs of different shapes, namely top-heavy (top, A and B), medium-heavy (middle row, C) and bottom-heavy (bottom, D and E) cascades (left, A, C and D) and pyramids (right, B and E).

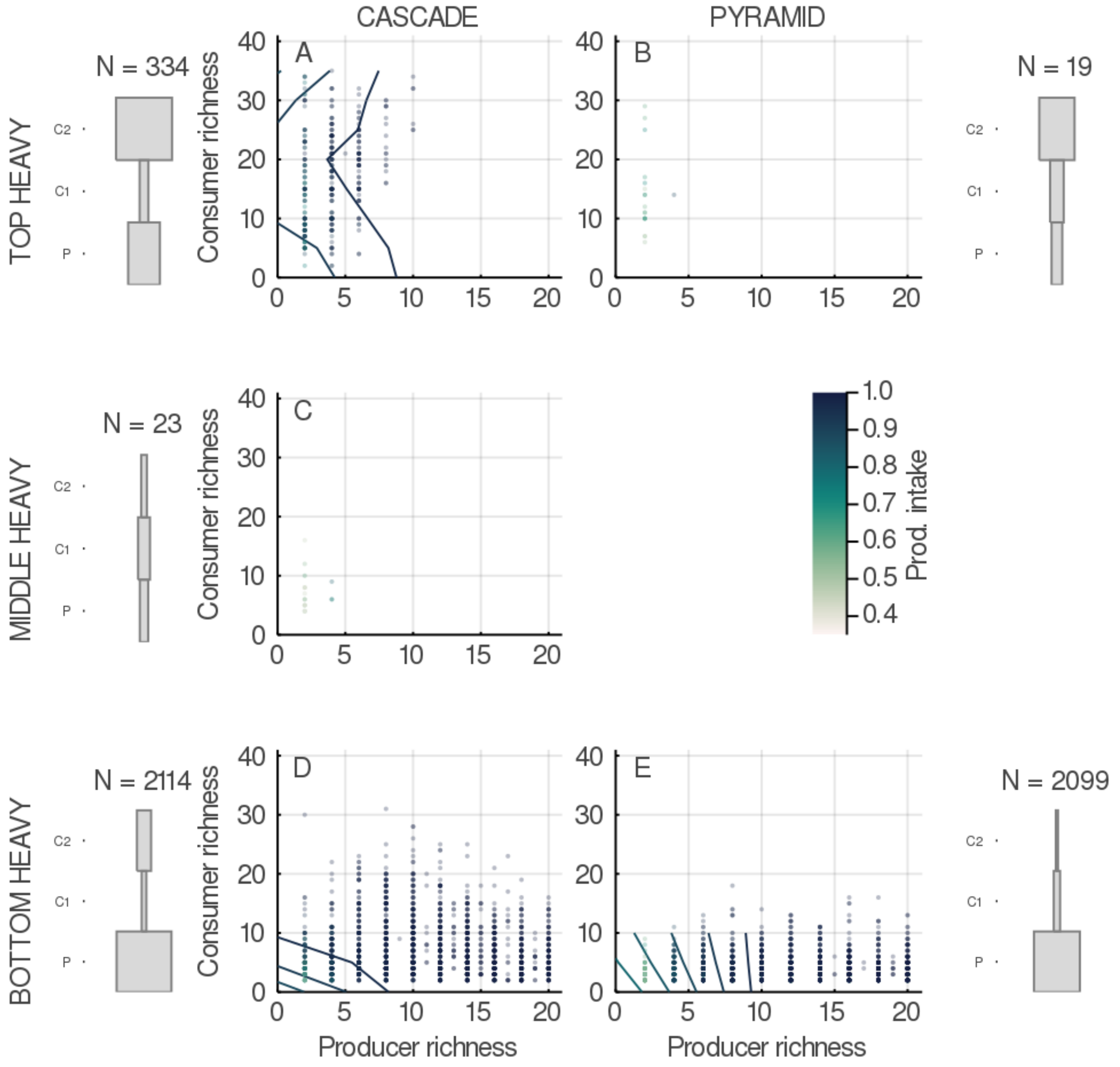

**Figure S 3** Diversity - total herbivory relationships in food webs. This figures shows the effect of producers and consumers richness on food webs total consumption by herbivores for food webs of different shapes, namely top-heavy (top, A and B), medium-heavy (middle row, C) and bottom-heavy (bottom, D and E) cascades (left, A, C and D) and pyramids (right, B and E).

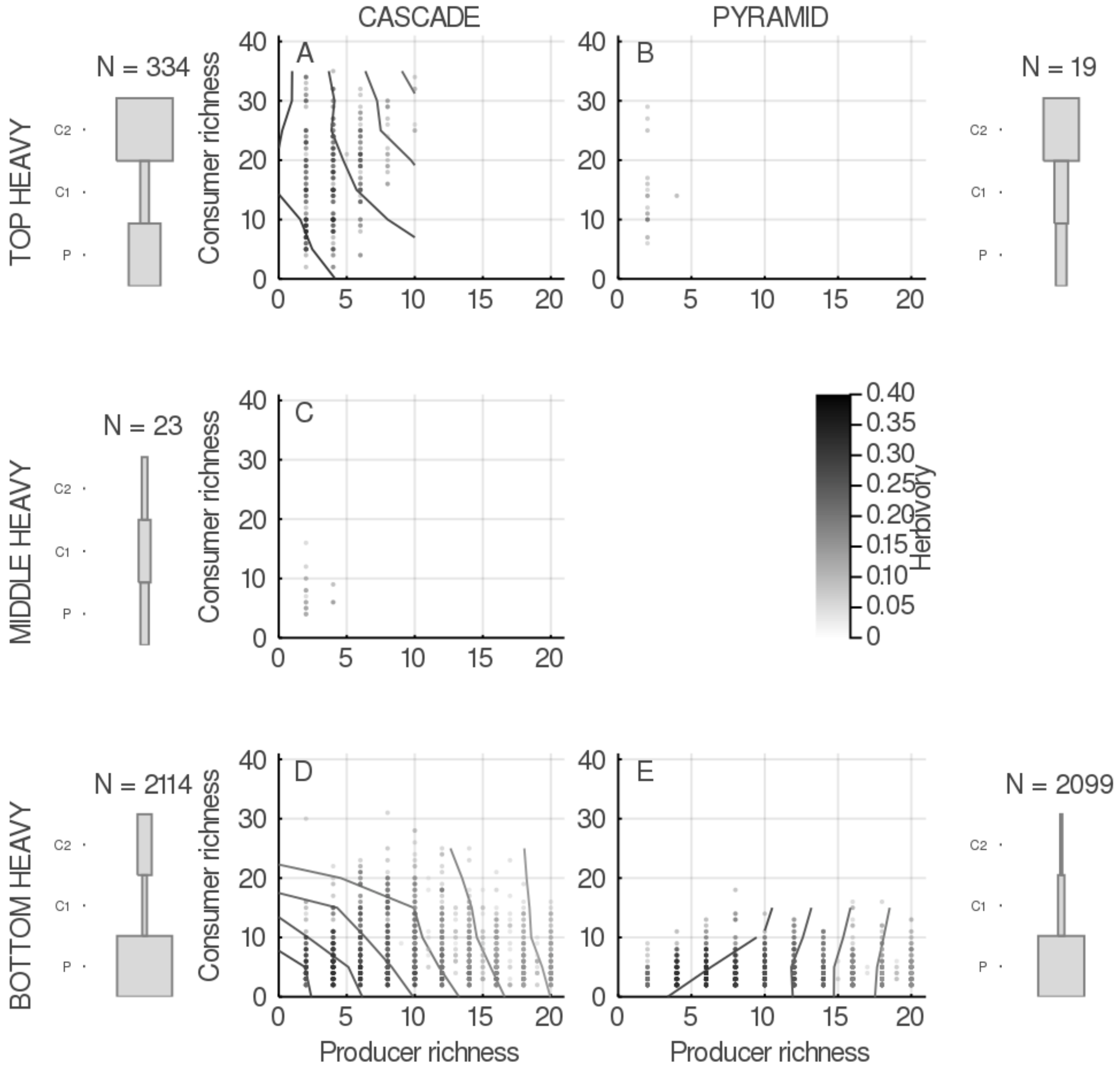

**Figure S 4** Diversity - total secondary consumption relationships in food webs. This figures shows the effect of producers and consumers richness on food webs total secondary consumption for food webs of different shapes, namely top-heavy (top, A and B), medium-heavy (middle row, C) and bottom-heavy (bottom, D and E) cascades (left, A, C and D) and pyramids (right, B and E).

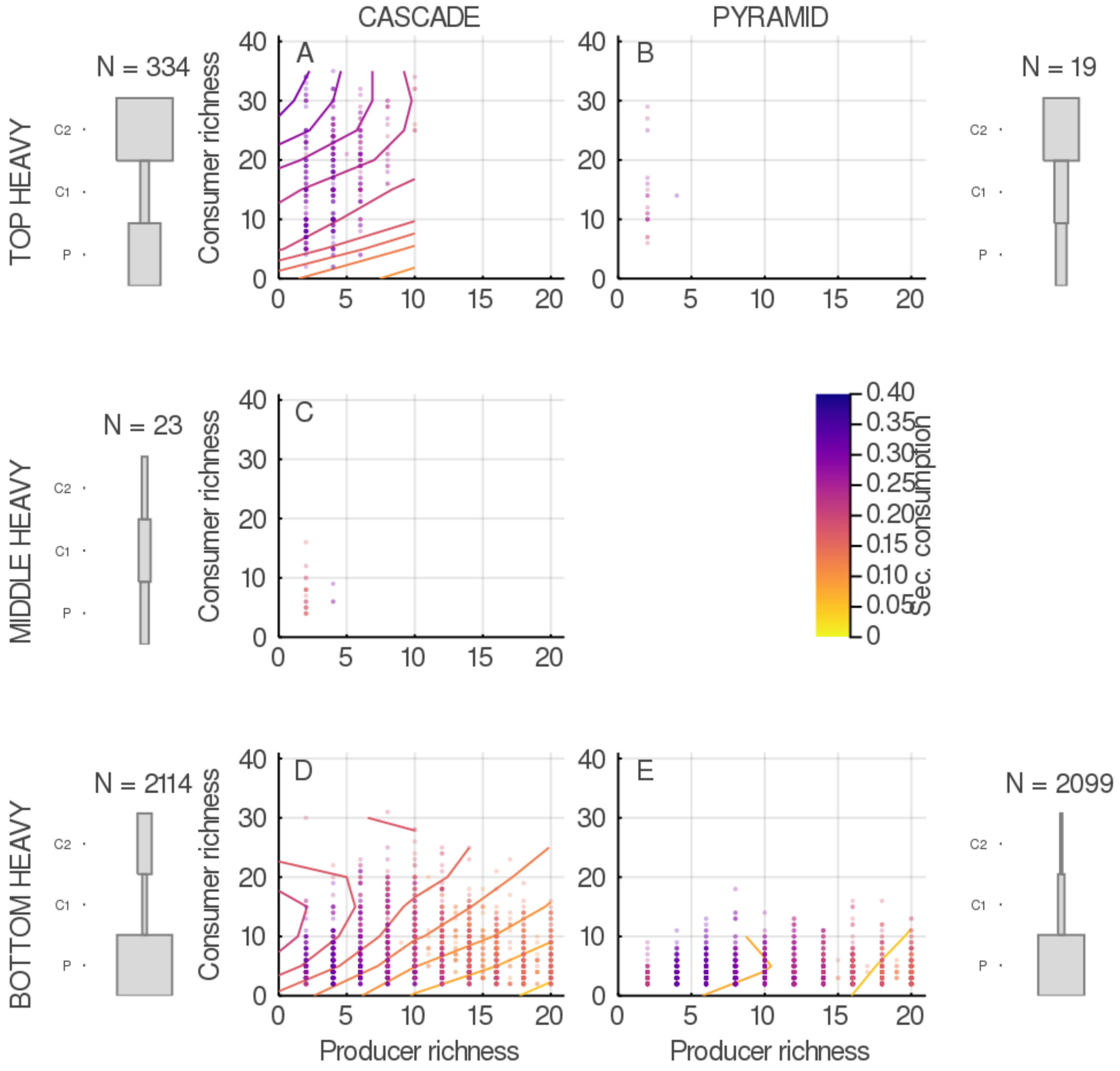
